# Heterotrimeric Collagen Helix with High Specificity of Assembly Results in a Rapid Rate of Folding

**DOI:** 10.1101/2024.05.21.595195

**Authors:** Carson C. Cole, Douglas R. Walker, Sarah A.H. Hulgan, Brett H. Pogostin, Joseph W.R. Swain, Mitchell D. Miller, Weijun Xu, Ryan Duella, Mikita Misiura, Xu Wang, Anatoly B. Kolomeisky, George N. Phillips, Jeffrey D. Hartgerink

**Affiliations:** Department of Chemistry, Rice University, Houston, Texas 77005, United States; Department of Bioengineering, Rice University, Houston, Texas 77005, United States; Department of Biosciences, Rice University, 6100 Main Street, Houston, Texas 77005, United States; Shared Equipment Authority, Rice University, Houston, Texas 77005, United States; Department of Chemical and Biomolecular Engineering, Rice University, Houston, Texas 77005, USA; Department of Physics and Astronomy, Rice University, Houston, Texas 77005, USA

## Abstract

The most abundant natural collagens form heterotrimeric triple helices. Synthetic mimics of collagen heterotrimers have been found to fold slowly, even compared to the already slow rates of homotrimeric helices. These prolonged folding rates are not understood and have not been studied. This work compares three heterotrimeric collagen mimics’ stabilities, specificities and folding rates. One of these was designed through a computational-assisted approach, resulting in a well-controlled composition and register, in addition to providing increased amino acid diversity and excellent specificity. The crystal structure of this heterotrimer elucidates the composition, register and geometry of pairwise cation-π and axial and lateral salt bridges. Complementary experimental methods of circular dichroism and NMR suggest the folding paradigm is frustrated by unproductive, competing heterotrimer species and these species must completely unwind to the monomeric state before refolding into the thermodynamically favored assembly. This collagen heterotrimer, which displays the best reported thermal specificity, was also found to fold much faster (hours vs days) than comparable, well-designed systems. The heterotrimeric collagen folding rate was observed to be both concentration and temperature-independent, suggesting a complex, multi-step mechanism. These results suggest heterotrimer folding kinetics are dominated by frustration of the energy landscape caused by competing triple helices.

## Introduction

Collagen-like proteins(CLPs) are found in all domains of life, including in prokaryotes and archaea. Their unifying structural feature is the polyproline type II triple helix. Prokaryotes utilize CLPs to bind to hosts during pathogenesis and for structural functions.^1^ In humans, collagen triple helices form the basis of more than fifty different types of CLPs.^2,3^ Fiber-forming collagens can be used as templates for biomineralization processes that give rise to rigid structures.^4^ The proteins C1q and SPA, which contain collagen-like triple helical domains, play essential roles in innate immunity.^5,6^ Most natural CLPs exist as heterotrimers, where strands in the triple helix are non-equivalent.

Synthetic, self-assembled triple helices of collagen mimetic peptides have served as models to study the structure and the stability of protein triple helices. Triple helices are the tertiary structure formed from three left-handed PPII monomers that fold into a supercoiled, right-handed helix.^7,8^ The common sequential motif of collagen is Xaa-Yaa-Gly where Xaa and Yaa are most frequently proline (P) and 4-hydroxyproline (O), respectively.^9^ The three peptide strands in a triple helix are offset by a single amino acid which differentiates the leading, middle and trailing peptide strands. Therefore, spatial registration differentiates an ABC register from ACB, BAC, BCA, CAB or CBA registers. Inter-strand pairwise interactions are termed axial and lateral^10^ and are essential for designing and controlling the register and composition of triple helices. Lys-Asp (KD), Arg-Tyr (RY) and Gln-Phe (QF) axial interactions have all been used to design triple helices and understand the chemical basis for collagen folding.^11–14^ Increasing amino acid diversity beyond these interactions is hampered by the fragility of the assemblies and is substantially more challenging when attempting to control the registration of a heterotrimeric triple helix.

While much of the literature has focused on understanding the collagen triple helix’s stability upon assembly, less attention has been paid to studying the folding pathway from free monomer to helix, especially in heterotrimers. The folding process for traditional collagens is believed to be driven by hydrophilic, and not hydrophobic, interactions.^15–17^ Collagen mimetic peptide folding starts with a nucleation event at the C-terminus.^18–20^ After nucleation, molecular dynamics and experimental studies with NMR have shown that *cis-trans* isomerization is the rate-limiting step for helical propagation down the axis.^21–23^ This complex mechanism has led to the hypothesis that the rate of collagen folding is effectively concentration-independent.^23^ For natural collagen folding, isomerases catalyze the *cis-trans* isomerization and once in the necessary *trans*-state, then folding can commence.^24–26^ Alternate theories suggest that the leading, middle and trailing strands’ folding propagate at different rates.^27^ Finally, it has also been proposed that the two strands can associate first, building a template for the third strand, or all three strands can simultaneously fold.^21^

The kinetics of folding have been modeled extensively in simple homotrimers containing (POG)*_n_*.^19^ The rate of folding for homotrimers has been found to vary according to the position along the axis, with C-terminal folding being second-order, while the central amino acids are best modeled as first-order.^27^ In addition, covalently tethered homodimers have been used to enhance the kinetic rate of assembly, similar to disulfide bond formation in natural collagens.^28,29^

Folding rates have yet to be studied in collagen heterotrimers for several reasons. First, the design of well-controlled, synthetic heterotrimers has only recently been accomplished.^30–32^ Second, quantifying the folding pathway and delineating if there are intermediates forming is challenging and requires the confirmation of both composition and register throughout the folding process.^33–35^ Finally, the heterotrimeric collagen helix is slow to form, days to months, and the pathway by which a competing heterotrimer can interchange with the most stable triple helix is unknown. This slow formation is contrary to the typical rapid formation of globular proteins such as lysozyme, which can fold on the order of microseconds to milliseconds,^36,37^ and triple helical homotrimer folding which can occur within minutes.^27^

Herein, we compare a series of synthetic ABC heterotrimeric collagen helices, Heterotrimer– 1 (HT1), Heterotrimer–2 (HT2) and Heterotrimer–3 (HT3), to investigate the kinetics and mechanism of folding. These were designed with the assistance of our scoring function, SCEPTTr.^32^ Of these, HT3 is notable because it possesses higher specificity over other compositions and registers in the folding landscape than any previously published triple helix. By using circular dichroism (CD) and NMR, specific folding to only the ABC register was confirmed. A successful crystal structure of HT3 reveals molecular-level insight into the geometry and interactions of cation-π and axial and lateral salt bridges. Folding rates were compared and HT3 was found to fold the most rapidly. The mechanism of the much faster folding HT3 is investigated in detail. Notably, when competing species are in solution, they hinder the folding process because they must first unwind into free monomer and can only then fold into the ABC register. This shows how an off-target folding event is difficult to remediate for heterotrimers. Therefore, we conclude that the energy landscape of competing species governs the overall rate at which the triple helix assembly occurs.

## Results and discussion

### Designing ABC Heterotrimers with better chemical diversity and specificity

The collagen scoring algorithm, SCEPTTr, was used to aid the design of three ABC-type heterotrimers, as shown in Table 1.^32^ In the current work, HT1-3 are named from lowest to highest melting temperature and specificity of their ABC-type assembly (see Table 2). HT2 is a previously published ABC-type heterotrimer and was designed with axial charge pair and cation-π interactions.^32^ The design of HT2 follows a straightforward geometric stabilization with the strongest pairwise charge pairing (K-D axial interactions) being used from strands A to B. Then B to C strands used K-E axial charge pair interactions. The C to A strands used R-F axial cation-π interactions. Additionally, prolines were incorporated at Yaa positions instead of hydroxyproline as a negative design to destabilize C homotrimers. Previous study^32^ of HT2 has shown that it forms the correct ABC register in solution and with relatively high specificity (T*_m_* = 39.5 °C, ΔT*_m_* = 18.5 °C).

**Table 1:**
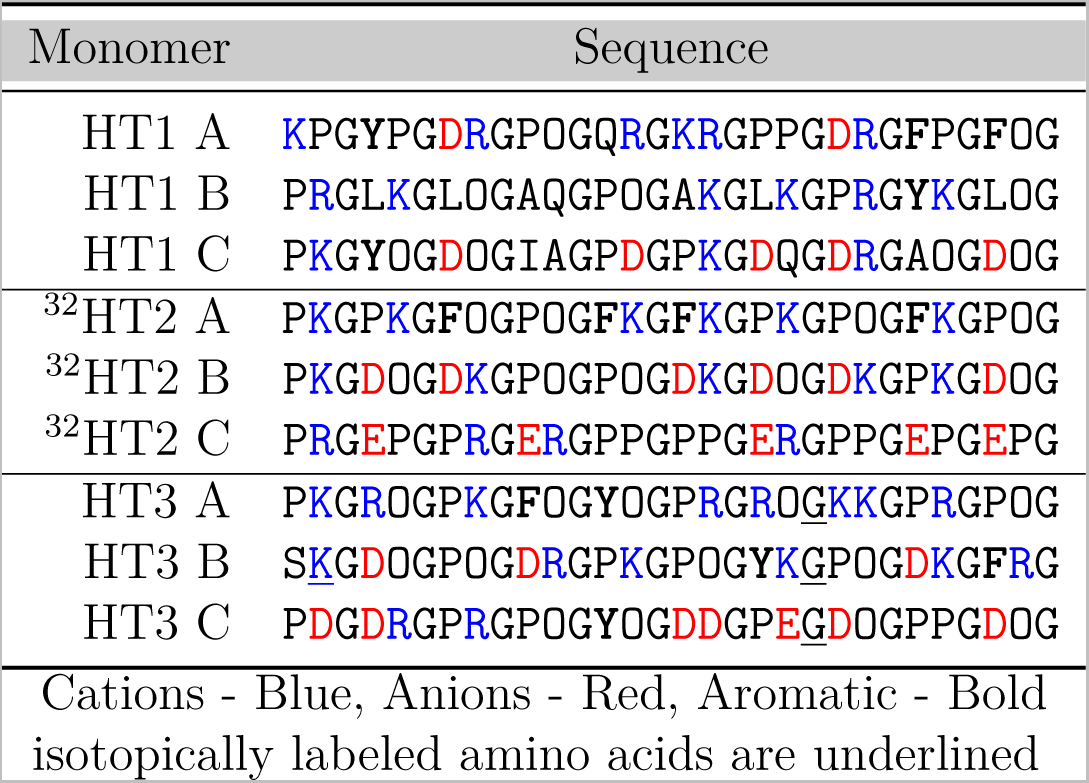
Heterotrimer Sequences of HT1-3.

**Table 2:** T*_m_* of Binary and Ternary Mixtures for Heterotrimers.

| Mixture | HT1 $T_m(^{\circ}\text{C})$ | HT2 $T_m(^{\circ}\text{C})$ | HT3 $T_m(^{\circ}\text{C})$ |
| --- | --- | --- | --- |
| ABC | 23.5 | 39.5 | 44.0 |
| AB | DNF | 20.0 | 17.5 |
| AC | DNF | 19.5 | 17.0 |
| BC | 12.5 | 21.0 | 14.0 |
| **Specificity** | **11.0** | **18.5** | **26.5** |

<sup>a</sup> DNF: Did not fold

Novel heterotrimers, HT1 and HT3, utilize these stabilizing design elements while also incorporating other stabilizing interactions and non-interacting, destabilizing amino acids. Hydrophobic amino acids such as leucine, isoleucine, alanine and Yaa-substituted prolines were incorporated into HT1’s sequence to limit the number of viable competing species (specificity: ΔT*_m_* = 11.0 °C, see Table 2). However, these amino acids also destabilize the ABC-type assembly (T*_m_* = 23.5 °C). HT3 utilizes non-interacting amino acids (B-Ser-1, B-Arg-29 and C-Pro-26) to promote the designed ABC register. In HT3’s A to B strands, there are three KD axial charge pairs and two axial cation-*π* interactions (RY and RF). The B to C strands interaction has three axial KD charge pairs. Finally, from the C to A strands there are three lateral salt bridges with two axial cation-π interactions. This design yielded not only a high stability of T*_m_* = 44.0 °C but also the best-reported specificity to date of ΔT*_m_* = 26.5 °C.

Seven mixtures of all components of the heterotrimers were studied with circular dichroism to determine melting temperatures of the various compositions (see Figure 2b, 2c, 2d). NMR was then used to confirm a distinct composition and ABC register in solution for HT3, which is presented in the supporting information (See Figures S14 and S15). Subsequent crystallization of HT3 (PDB ID 8TW0) also confirmed the correctly predicted composition and register for the triple helix in addition to confirmation of the theorized geometries of the pairwise interactions employed.

**Figure 1:**
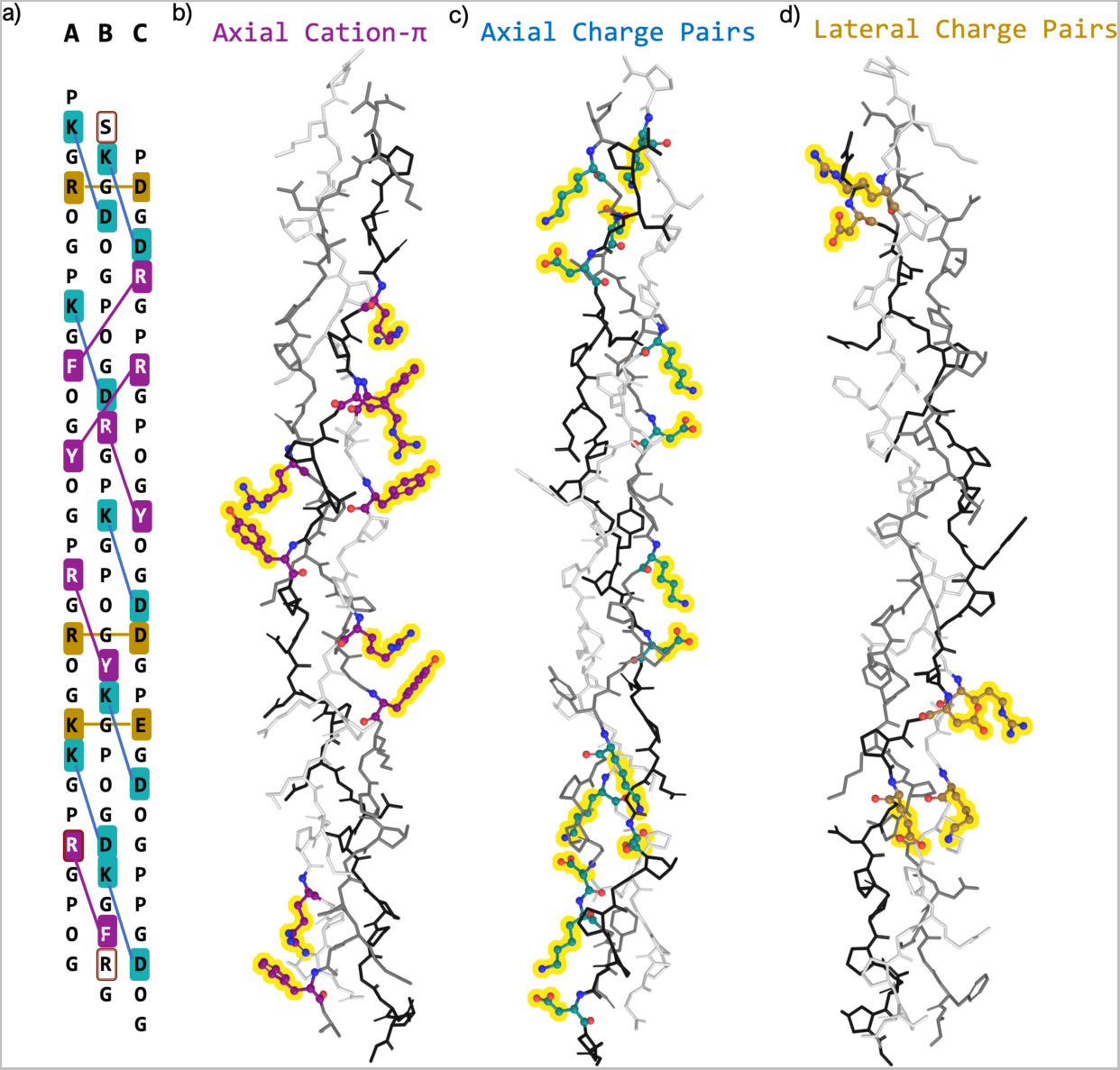
Sequence and Crystal Structure of HT3. a) The sequence of HT3 shows a variety of pairwise interactions with axial (blue) and lateral charge pairs (gold), axial cation-π (purple) interactions, and non-interacting (brown box) residues b) Crystal structure of HT3 (PDB ID: 8TW0) shows the cation-π geometries c) Highlights of the charge pairs that create stabilizing axial salt bridges d) Lateral salt bridge geometries from the same PDB.

**Figure 2:**
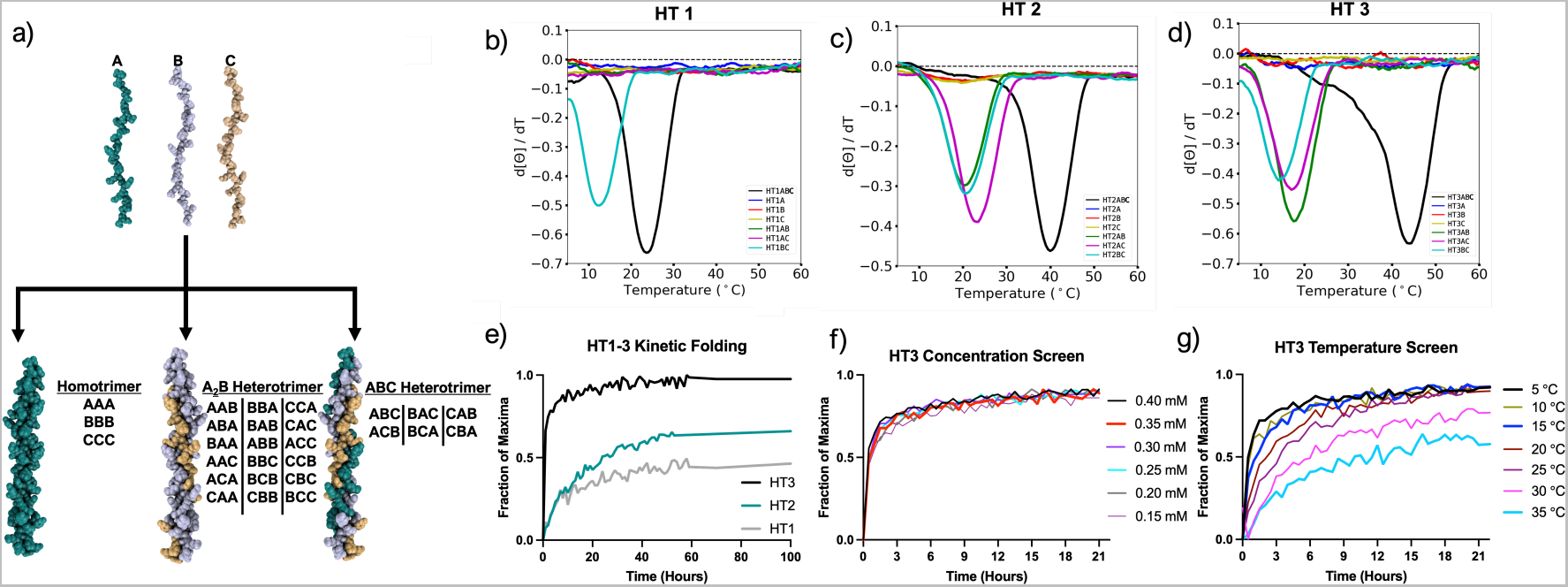
Monitoring Heterotrimer Folding with Circular Dichroism. a) The twenty-seven possible combinations of a triple helix b) HT1 melt derivative curves c) HT2 melt derivative curves d) HT3 melt derivative curves e) The ABC-type heterotrimer of each HT1-3 and their second most stable helix f) CD of the HT3 ABC solution by varying the total peptide concentration folding was performed at 5 °C g) Temperature screen of HT3 folding at 0.3 mM.

### Crystal Structure of HT3 at 1.53 Å resolution

This is the first reported crystal structure of cation-π interaction geometries at the molecular level for any collagen mimetic peptide (CMP) triple helices. In the crystal structure, pairwise axial cation-π interactions have an average interaction distance of 4.1 Å between the center of the aromatic ring of Tyr (Y) or Phe (F) with the *δ*-guanidinium cation’s central carbon of Arg (R) (see Figure 1b). The two theorized geometries of planar and T-shaped were also observed for the pairwise interactions. The KD axial charge pairs, highlighted in Figure 1c, showed the closest distance for any pairwise interaction with the average distance of 3.6 Å and near ideal N — O hydrogen bonding distance of 2.7 Å, as measured between the *’*-nitrogen of Lys (K) to the *γ*-carbon of Asp (D). This once again demonstrates KD axial interaction’s robustness in de novo design. Reverse, lateral charge pairs were measured from the *’*-nitrogen of Lys (K) or the *δ*-guanidinium cation of Arg to the *γ*-carbon of Asp (D) or the *”*-carbon of Glu (E) and had an average interaction distance of 3.9 Å and a corresponding hydrogen bonding distance of 2.8 Å (see Figure 1d). All interactions are summarized in the supporting information’s Table S1.

### Determining Folding Kinetics with Circular Dichroism

With HT3’s register and composition confirmed, we further investigated how ABC-type heterotrimers assemble. The task of selectively folding an ABC register and composition from 27 different possible intermediates (Fig. 2a) is non-trivial. Collagen mimetic heterotrimers have been known to assemble slowly, usually on the order of weeks to months. We discovered that HT3 assembles to more than 91% in a single day which allows us to address the previously inaccessible question of triple helical heterotrimer folding kinetics. We performed time-lapse CD experiments on HT3 to study folding kinetics and performed similar experiments on HT1 and HT2 as a point of comparison. For HT1-3, the heterotrimer’s mixture indicates that the ABC heterotrimer has the highest thermal stability. Of the three, HT1 has the lowest specificity (11.0 °C), but also the fewest observable competing helices (Fig. 2b). HT2 (Fig. 2c) exhibits an excellent specificity (ΔT*_m_*) of 18.5 °C but with more observable stable competing species. HT3 has the highest stability (T*_m_* = 44.0 °C) and specificity (ΔT*_m_* = 26.5 °C) as demonstrated by CD in Figure 2d. All of the identifiable folded species for HT1-3 are listed in Table 2.

The polyproline II (PPII) secondary structure yields a maximum CD signal at around 225 nm and the mean residual elipticity (MRE) is correlated to triple helix assembly. The maximum MRE for each heterotrimer was determined by measuring the CD of the folded ABC mixture after one month of folding at 4 °C. For all experiments, each sample’s observed MRE was then divided by this maximum value at 5 °C to yield a ”fraction of maxima.” Monitoring the triple helix formation in the case of ABC-type heterotrimer formation does not guarantee that solely the ABC-type register is observed; however, we did observe relative folding rates for HT1, HT2, and HT3. The folding of HT3 nearly reached completion in forty-eight hours (Figure 2e). HT1 and HT2, because of their slower folding nature, were monitored for thirty days. The maximum signal was then used to normalize the kinetic plots and are presented in Table 3. To fold to 50% of the observed fraction it took HT1 and HT2 147 hours and 27 hours, respectively. This is considerably slower than HT3, which took a mere 0.50 hours.

**Table 3:**
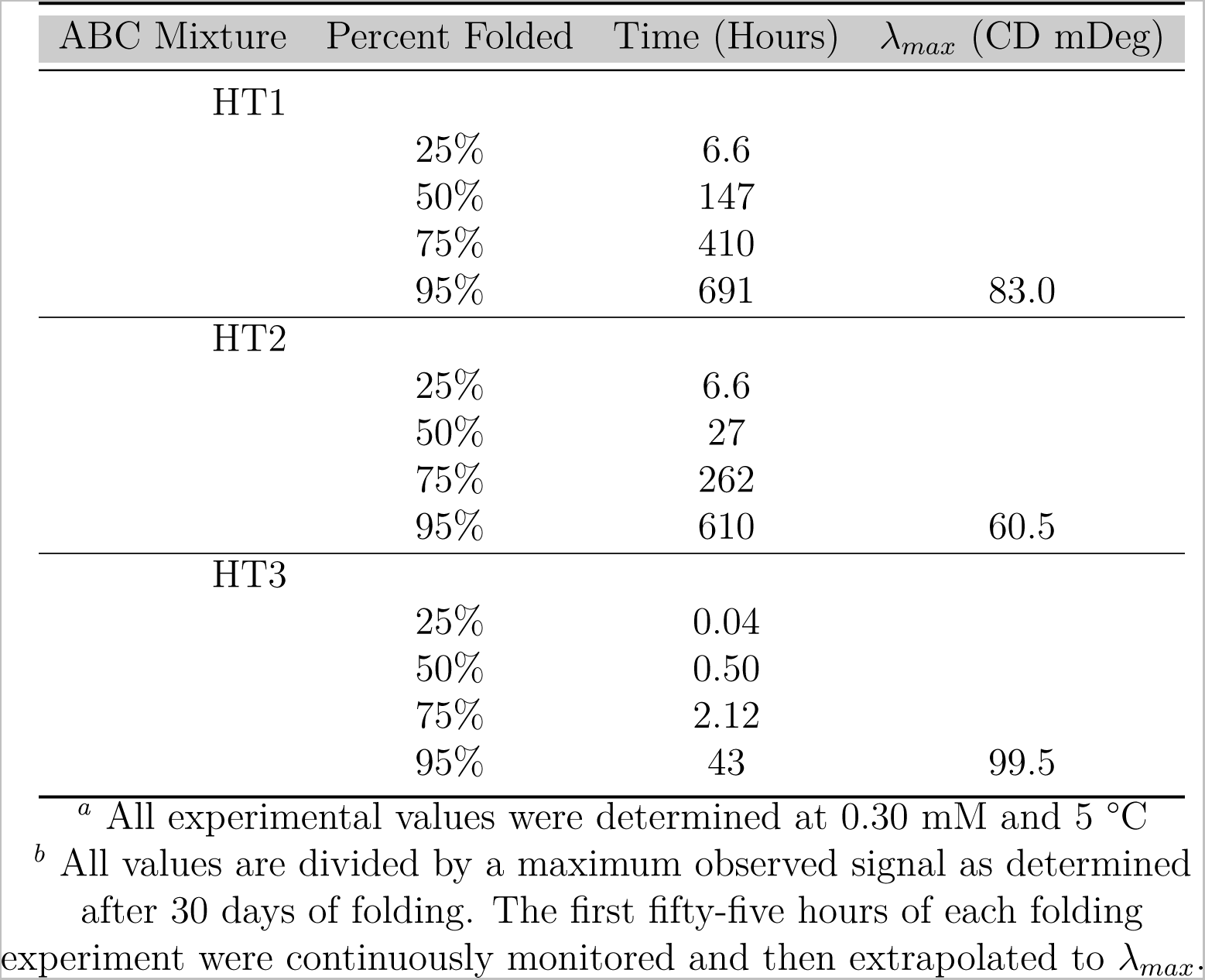
Circular Dichroism Kinetics of HT1, HT2 and HT3.

Monitoring the formation of HT3 at various concentrations and temperatures, without denaturant, yields further insight into heterotrimer folding. In Figure 2f, we see that the folding kinetics are concentration independent, in the range of 150 *µ*M-400*µ*M. Using an exponential plateau function we also calculated relative rate constants which showed that that the formation for each sample was not statistically significant (see supporting information Tables S3 and S4). The concentration independence supports that the nucleation step in collagen folding is not governing the folding paradigm. Therefore, similar to homotrimer folding, heterotrimer folding is limited in its rate by isomerization.^26^

A temperature variability study shown in Figure 2g appeared to show a decrease in kinetic folding rate as temperature increased (see supporting information Tables S5 and S6). By more closely examining thermal melting derivative curves of HT3 in Figure 2d we observed a loss in CD signal starting at 20 °C. Therefore, we correlated the maxima for each temperature (supporting information Figure S11a), and used that maxima to plot the fraction folded for each temperature. At temperatures below 20 °C the heterotrimer folding rate was consistent and agrees with previous studies which were performed on homotrimers.^23^ However, at temperatures greater than 20 °C the systems kinetic folding rate was perturbed due to increasing denaturation.^38^

### Heterotrimer Folding Kinetics with NMR

The CD experiments described above provided relative folding rates. Nevertheless, there is no indication if there is a mixture of compositions and/or registers or if a single ABC-type heterotrimer is contributing to the increase in signal (see supporting information Figure S13b). By using NMR, folding specific to particular compositions and registers could be investigated. The formation of HT3 was observed with heteronuclear single quantum coherence (HSQC) NMR by labeling strands A, B and C with a ^15^N-amide glycine at the 21st position (Table 1). Strand B also incorporated a ^15^N-^13^C label at the Lys-2 position. We correlated each peak to the free monomer and assembled a ABC heterotrimer (see supporting information Figure S15). These monomer and trimer peaks are in equilibrium with each other in solution.^27^ By heating the sample above the melting temperature of HT3 (T*_m_* = 44.0 °C) to 50 °C, A, B and C peptides are all unfolded. Assembly was then monitored at 5 °C and carried out in triplicate.

Figure 3b plots the time course of HSQC of the A/B/C strands upon assembling into the ABC heterotrimer. The intensity of each assembly strand is statistically similar, and the total folding event for the trimer is shown in black (see supporting information Figure S21c). Throughout the time course of each experiment, no transient signal was observed other than the monomer and trimer peaks. We observed that as the ABC heterotrimer formed, the monomer signals decreased, as seen in 3c. By using Bruker Dynamic Center the first-order fit was used to determine k_1_ and these data are summarized in supporting information’s Table S7. The A-strand formed at a rate of 0.019 *±* 0.003 min^-1^. For the B and C strands each formed at a rate of 0.021 *±* 0.002 min^-1^ and 0.021 *±* 0.004 min^-1^, respectively. Which is considerably slower than the known homotrimer folding k of 1.38 min^-1^.^39^ In Table 8 of the supporting information, we also demonstrate the same trend of decreased folding rate with increased temperature which is consistent with the CD data and denaturation.

**Figure 3:**
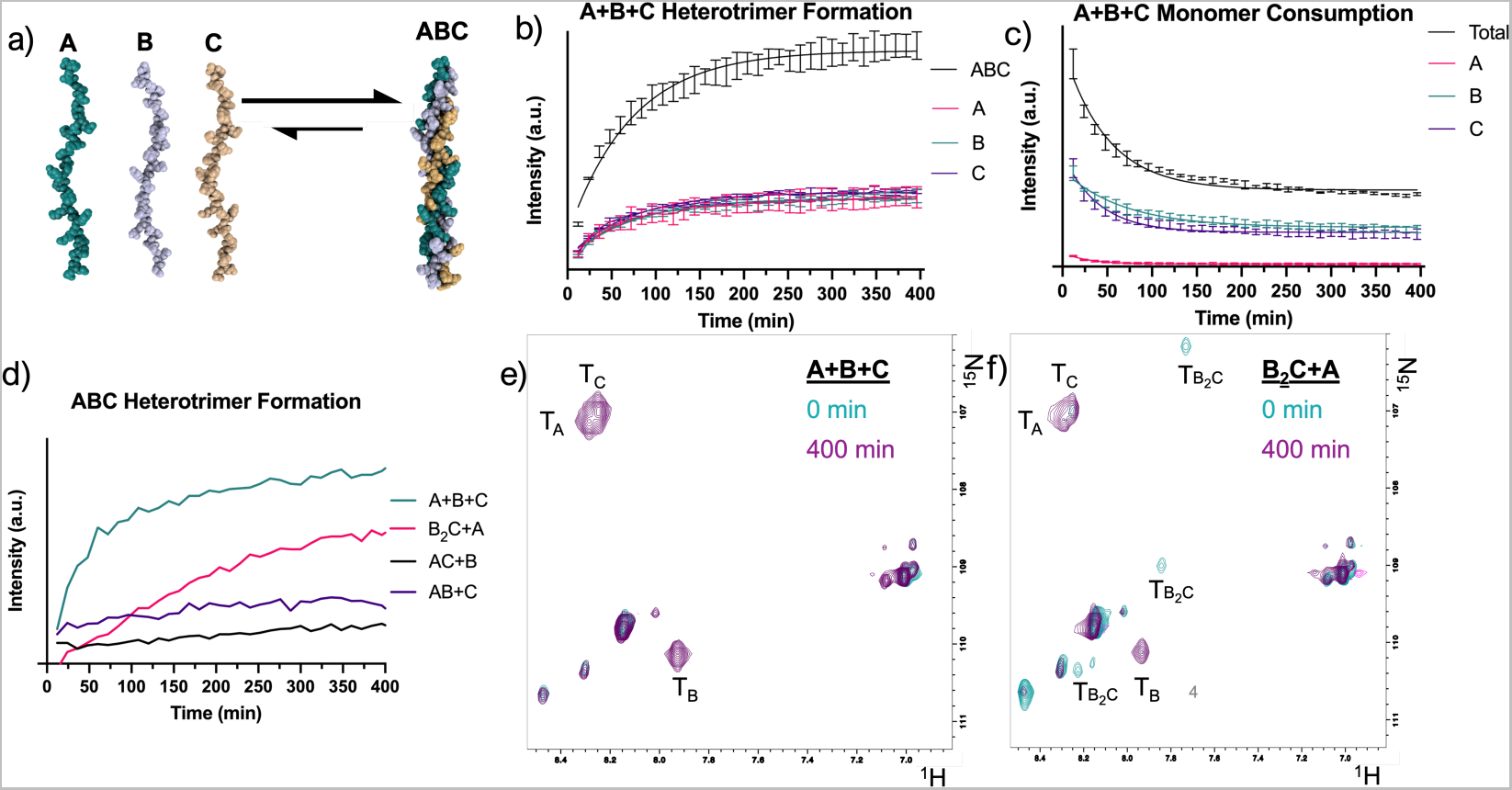
Free Monomer Folding into HT3. a) Illustration of A, B, and C monomers assembling into HT3 b) The kinetic folding curve of each individual peak of HT3 and their total intensity (black) c) Monomer consumption of free monomers and their total decrease (black) d) Discrepancies in the folding curves depending on the starting solution e) HSQC spectra from the initial and final time-points of the heated HT3 system with the free monomers assembling f) The same HSQC spectra but with a B_2_C heterotrimer in solution. The B_2_C heterotrimer peaks disappear after HT3 assembles. Spectra were obtained at 5°C and fit with an exponential plateau function. All confidence intervals were set to 95% as determined by weighted fit for n = 3.

In heterotrimer folding, the folding pathway can be frustrated by competing species that are preformed. To investigate how preformed heterotrimer may change the kinetic rate of folding, we carried out an additional experiment where samples with two components of the ABC system were annealed and folded for one week at 4 °C. The AB, AC and BC mixtures form a number of competing trimers with melting temperatures of 17.5, 17.0 and 14.0 °C, respectively. Further experimentation confirmed a B_2_C heterotrimer arose from the BC mixture (supporting information Figure S19 and Table S7). The missing peptide strand was then added to the preformed heterotrimer solution. Evolution of the system was then monitored as before by HSQC.

As summarized in Figure 3d, we observed that the initial heterotrimers being in solution perturbed the rate of folding as compared to the mixture that contained only monomers. The B_2_C+A solution (pink line) was the most readily quantifiable amongst the pre-folded mixtures and also has the lowest stability (T*_m_* = 14.0 °C). The k_1_ of each sample is summarized in Table S7 of the supporting information. Additionally, when comparing Figure 3e and Figure 3f the B_2_C trimer peaks were no longer detectable after 400 minutes. These data support our supposition that formed heterotrimers must unfold before they can refold to a more thermally stable triple helix.

The general mechanism for heterotrimer formation with a previously formed heterotrimer (B_2_C) is outlined in Figure 4a and outlines a complex equilibrium-based assembly mechanism. Through HSQC experiments in Figure 4b, it was clear that as the ABC heterotrimer formed (black), the B_2_C heterotrimer’s intensity decreased (light blue). Figure 4c shows that as A peptide is consumed, the B_2_C heterotrimer also disappears. Thus as the free B and C peptide interacts with the free A peptide to form the ABC assembly, the equilibrium shift drives the B_2_C heterotrimer to unwind into the free monomers. Despite the experimental temperature being below the thermal stability of B_2_C, the folding of HT3 still occurs, further supporting an equilibrium in which full unwinding occurs. In contrast to a concerted step where a single strand would be replaced directly by another strand.

**Figure 4:**
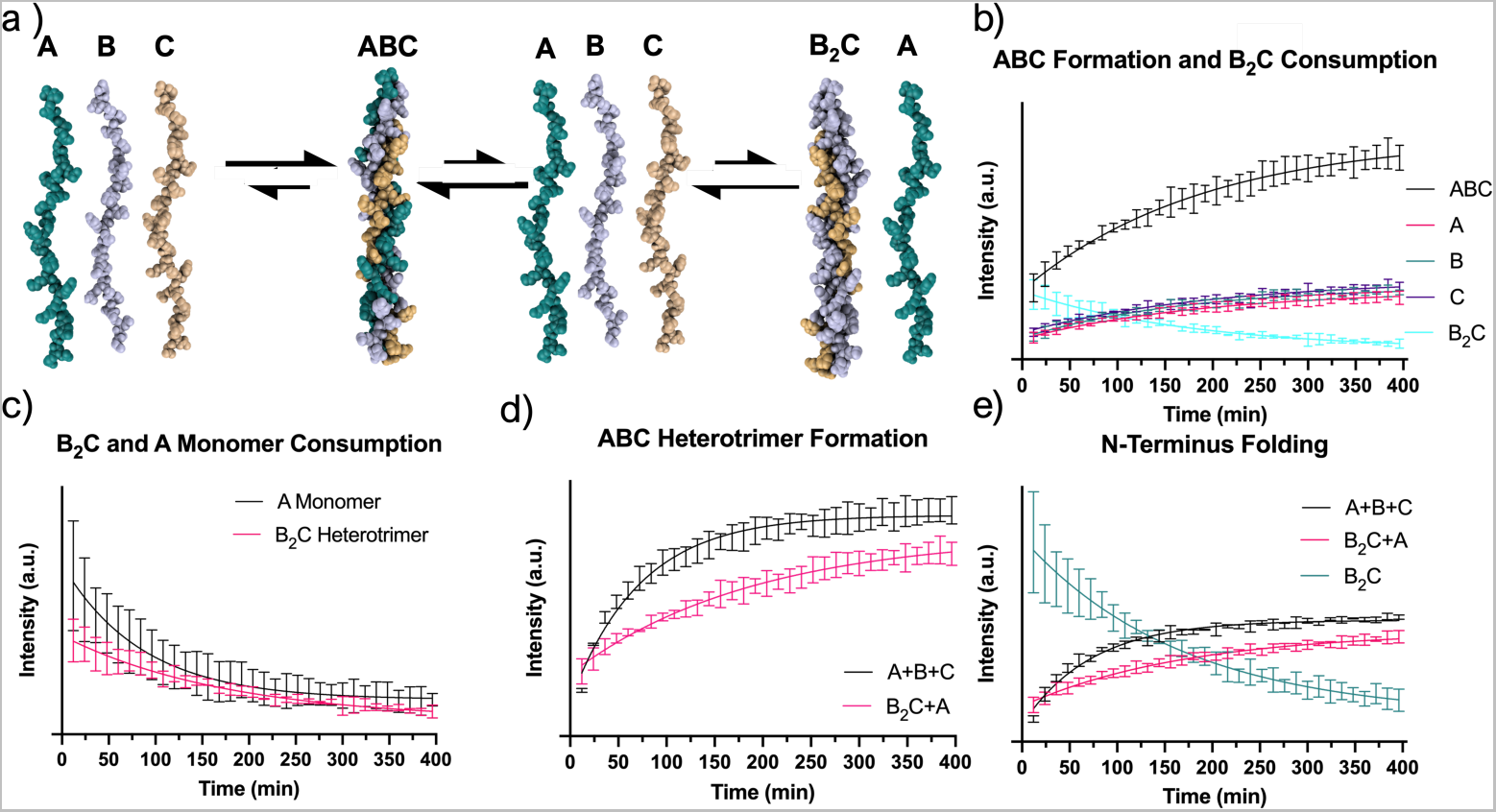
B_2_C Folding Impacts HT3 Formation. a) Illustration of the two general pathways of HT3 formation with a preformed heterotrimer b) The formation of HT3 (black) is inversely proportional to the loss of the competing B_2_C heterotrimer c) B_2_C’s heterotrimer disappearance correlates to the disappearance of the A monomer d) The triplicate sets of HT3 formation from free monomer (A+B+C) and the B_2_C solution (BC+A) e) The N-terminus folding of each mixture from the 2-Lys in monomer B for each triplicate set. Spectra were obtained at 5°C and fit with an exponential plateau function. All confidence intervals were set to 95% as determined by weighted fit for n = 3.

Figure 4d plots the formation of the ABC assembly from the two studies. The rate of formation from free monomers was more rapid (A: k_1_ = 0.019 *±* 0.003 min^-1^ B: 0.021 *±* 0.002 min^-1^ C: 0.021 *±* 0.004 min^-1^ ) when compared to the solution comprising B_2_C in a preformed state (A: k_1_ = 0.005 *±* 0.001 min^-1^ B: 0.0.005 *±* 0.001*^≠^*^1^ min^-1^ C: 0.005*±* 0.001 min^-1^). In Figure 4e the same trend was observed for the N-terminal folding in each system. Where the N-terminus folding was experimentally observed to form at the same rate as the other Gly-21 HSQC of the A, B and C strands (See supporting information Table S8). This is conclusively demonstrating that the unwinding event of the competing heterotrimer species is necessary for the formation of the ABC assembly and demonstrates the path-dependent folding for heterotrimeric collagens.

The equilibrium dependency of heterotrimers demonstrates that the off-target association of monomer strands in solution greatly impacts the rate of triple helix assembly and explains why, in the cases of HT1 and HT2, months have been required for heterotrimers to reach maximal signals in CD. Natural collagens fold more readily by forming disulfide bridges, possessing pro-peptide regions, and through the assistance of chaperonins, all of which help to guide the formation of the appropriate assembly and can subsequently be cleaved off.^3,27^

The design of HT3 with computational aid generated a heterotrimer with the highest specificity to date. Further the rate of folding of this heterotrimeric helix also resulted in a folding rate that was rapid enough to be readily observed experimentally. Due to the specificity of HT3, it is able to avoid kinetic traps and achieve the targeted and thermodynamically stable, ABC register. This next-generation design expands our understanding of cation-π interactions, incorporates non-interacting amino acids, and varies pairwise interaction propensities between each strand. This complex de novo design success shows how a detailed supramolecular understanding of pairwise interactions can motivate the investigation of biologically relevant collagen sequences and the next generation of collagen mimetic biomaterials.

## Experimental

### X-ray Crystallography

#### HT3 Crystallization

The assembled HT3 heterotrimer was diluted to 20 mg/mL in MES buffer (10 mM, pH = 6.1). HT3 was then annealed at 85 °C for 15 minutes before being cooled to room temperature. A diluted aliquot of HT3 was then confirmed to fold with circular dichroism. Subsequently, high-throughput crystal screening was performed using a Mosquito LCP robot (SPT LabTech, Melbourn, UK) to set up vapor diffusion sitting drop experiments with commercial screens from Hampton Research (Aliso Viejo, CA), IndexHT and PegRx-HT, and Rigaku Reagents (The Woodlands, TX), WizardHT screens 1-2 and 3-4. Drops formed by mixing 200 nL peptide and 200 nL reservoir solution. These were equilibrated against a reservoir of 200 *µ*L at 20 °C. Of the initial 384 conditions screened, over 35 conditions produced crystals hits within one week and more than 25 conditions had crystals large enough to mount.

#### Crystal data collection and structure refinement

A crystal looped from the Index-G8 condition (0.2 M Ammonium acetate, 0.1 M HEPES pH 7.5, 25% (w/v) Polyethylene glycol 3,350) was flash-cooled in a 100 K nitrogen stream (Oxford Cryosystems, cryostream 800). Data were collected to 1.53 Å using a Rigaku (The Woodlands, TX) XtaLAB Synergy-S-DW diffractometer equipped with a HyPix-Arc 150° Hybrid Photon Counting (HPC) detector and Cu microfocus sealed X-ray source. Data were integrated, scaled and merged within the CrysAllis Pro software package.^40^ Phasing was accomplished with Phaser^41^ and shelxe^42^ within the Arcimboldo pipeline,^43^ which placed 3 copies of a blunt-end starting model derived from PDB id 3t4f^44^ (residues A:8-16, B:7-15, C:6-14 with Lys and Glu residues truncated at the C*^—^* position). Shelxe auto-traced 75 residues with a final CC of 41%. ARP/wARP^45^ was used for building side chains and built

76 residues. The HT3 model was inspected and rebuilt in Coot^46^ and refined with phenix.^47^ PyMOL was used for structure visualization and analysis. The structure was visualized using a collaborative 3-D graphics system.^48^ Structural biology software applications used in this project were compiled and configured by SBGrid.^49^ Data processing and refinement statistics are summarized in Table S2. Atomic coordinates of HT3 (PDB ID:8TW0) crystal structures have been deposited with the Protein Data Bank. The authors declare that the data supporting the findings of this study are available within the paper and its Supplementary Information files. Should any raw data files be needed in another format they are available from the corresponding author upon reasonable request.

### NMR

#### Structural Studies

All NMR experiments were carried out on a Bruker NEO 600 MHz NMR spectrometer equipped with a 5-mm TCI cryoprobe. Isotope-labeled collagen peptide was resuspended to 2.7 mM in an NMR buffer containing 50 mM phosphate, 50 mM KCl, 10 *µ*M trimethylsilyl propanoic acid (TSP), and 90%H2O/10% D2O (pH 6.5). The nitrogen carrier frequency was set at 117 ppm for the ^1^H, ^1^H, ^15^N filtered 2D NOESY-HSQC for all experiments. A spectral width of 25 ppm was used in the nitrogen dimension for all experiments. The proton carrier frequency was set at the water signal for all experiments and a spectral width of 25 ppm was used in all HSQC experiments. In the NOESY-HSQC, the first and second proton dimension values were identical to those used in the HSQC experiments. HT3 experiments were collected at 5 °C, 15 °C, 25 °C, 30 °C and 35 °C. For the HSQC experiments, 32 dummy scans were run before each experiment, then 8 scans were for each transient in the experiments. For all HSQC experiments, 256 transients were collected in the nitrogen dimension. For the NOESY-HSQC experiments, 2048 x 1 x 700 complex data points, with 8 scans per data point, and 128 dummy scans were run. 3D 15N-edited NOESY-HSQC was performed with a mixing time of 100 ms at 25 °C. Chemical shift referencing: 1H chemical shifts were referenced to TSP and 15N chemical shifts were referenced indirectly according to the gyromagnetic ratio.^50^ NMR data were processed using Bruker TopSpin software. Pymol and DLPacker were used to render energy-minimized structures for determining the correct register.^51,52^

#### Kinetics

To quantify the kinetics of HT3 formation with NMR HSQC experiments were optimized at 5 °C. Briefly, three independent solutions were prepared to have a final total peptide concentration of 2.8 mM in a 10% solution of D_2_O with TMSP. For the formation of HT3 from free monomers (Sample: A+B+C), the sample was loaded and parameterized at 5 °C, then the sample was heated for one hour at 55 °C. After one hour, the sample was cooled to 5 °C and HSQC spectra was immediately collected with the same parameters as above. For all HSQC experiments, 256 transients were collected in the nitrogen dimension. Samples were collected for 400 min, each scan took 10 minutes and 29 seconds. The next scan started 90 seconds later, giving a twelve-minute step between each data point. For the competing heterotrimer studies, such as BC+A, the heterotrimer and monomer strands were annealed and folded for one week. After folding the O1P was determined on a sample prepared in parallel and then both samples were taken from 5 °C, combined, and then placed into the magnet. After shimming and the loading procedure sample signal was collected starting at fifteen minutes. The same HSQC experiment was conducted with a step of twelve minutes for each run. Data analysis was used to correlate spectra peaks, to identify compositions and then were plotted in GraphPad Prism software. All equations are reported and rationalized in the supporting information.

## Conclusions

Here, a collagen triple helix, termed HT3, was computationally designed to have controlled specificity and register by utilizing a variety of charge pairs, cation-π interactions and non-interacting amino acids which led to the most specific heterotrimer in literature. By characterizing this triple helix structurally with crystallization, pairwise interaction geometries and distances were able to be visualized and quantified. NMR analysis concluded that the system folded into a distinct heterotrimer in solution, and circular dichroism studies suggested that specificity and lack of competing species can drive an increase in heterotrimer assembly rate. By CD the system was determined to be concentration independent during folding. The assembly mechanism and rate of HT3 formation were then monitored with NMR. Mechanistically, collagen heterotrimer folding is motivated by varying equilibria in solution and is path dependent. This work informs and demonstrates the biophysical phenomena that belie the assembly process of the body’s most abundant protein, collagen.

## Author Contributions

Carson C. Cole: conceptualization, data curation, methodology, investigation, writing. Douglas R. Walker: conceptualization, methodology, writing, investigation. Sarah A.H. Hulgan: methodology, investigation. Brett H. Pogostin: investigation, data curation. Joseph W.R. Swain: investigation, data curation. Mitchell D. Miller: methodology (crystallography), investigation. Weijun Xu: methodology (crystallography), investigation. Ryan Duella: methodology, investigation. Mikita Misiura: investigation, methodology. Xu Wang: methodology (NMR), investigation. Anatoly B. Kolomeisky: methodology (kinetics), investigation. George N. Phillips, Jr: methodology (crystallography), investigation. Jeffrey D. Hartgerink: conceptualization, investigation, editing, writing, supervision, funding acquisition.

## Acknowledgement

This work was funded in part by the National Science Foundation (CHE 2203937), National Science Foundation BioXFEL STC (1231306), Robert A. Welch Foundation Grants (C-2118, C-2141). and the National Science Foundation Graduate Research Fellowship Program (Grant No. 1842494). Any opinions, findings, conclusions, or recommendations expressed in this material are those of the author(s) and do not necessarily reflect the views of the National Science Foundation. This work was done in part using resources of the Shared Equipment Authority at Rice University.

## Supporting Information Available

Full peptide characterization including UPLC, mass spectrometer data, CD melt curves; further NMR characterization at various temperatures; other kinetic plots.

## Supporting Information Table of Contents Methods Continued

### Peptide Synthesis

Peptides were synthesized using standard Fmoc-protected amino acids on a low-loading rink amide MBHA resin to leave final peptides with C-termini amidation. 25% (v/v) piperidine in dimethylformamide (DMF) was used for deprotection. Coupling steps were done with 2-(1H-7-azabnzotriazol-1yl)-1,1,3,3-tetramethyl uranium hexafluorophosphate methanaminium (HATU) and diisopropylethylamine (DiEa) in DMF at a 1:4:4:6 equivalents of resin:amino acids:HATU: DiEA. The N-terminus was acetylated with a double addition of excess acetic anhydride and DiEA in dichloromethane (DCM). Peptides were cleaved from the resin with 10% v/v scavengers in trifluoroacetic acid (TFa). Scavengers used include triisopropylsilane, H_2_O, anisole, and ethanedithiol. All peptides were synthesized with a ^15^N isotopically enriched glycine at the 21st position and HT3’s B monomer contained an enriched lysine in the 2nd position. These positions are underlined and bolded in Figure S**??**A below. After cleavage, the TFA mixture was moved to a round bottom vessel. TFA was removed from the mixture by evaporation under nitrogen pressure. Cold diethyl ether precipitated peptide from the remaining mixture of scavengers. This was centrifuged and the ether was decanted and then repeated. Peptides were dissolved in H_2_O and filtered. They were then purified by reverse phase high-performance liquid chromatography (HPLc) with water and acetonitrile at a gradient of 0.7%/min on a 19 x 250 mm C-18 column. The water and acetonitrile both were treated with 0.05% TFA. Samples were rotovapped, frozen, and lyophilized. A matrix-assisted laser desorption/ionization mass spectrometer (MALDI-MS) confirmed the correct peptide mass.

### Circular Dichroism

#### Structural Studies

Peptides were dissolved in Milli-Q water and concentration was determined by mass or by using a ThermoFisher Nanodrop Spectrophotometer. Samples were diluted to 3 mM peptide in 10 mM phosphate bu↵er at pH 7.0. All mixed peptide samples were prepared to 3 mM total peptide. Samples were preheated at 85 °C for 15 minutes, cooled to room temperature, and refrigerated for a week before running CD. Polyproline type II (PPII) helix formation, which is correlated to triple helix formation, was measured with a Jasco J-810 spectropolarimeter (Tokyo, Japan) equipped with a Peltier temperature-controlled stage. An aliquot of 200 *µ*L of the 0.3 mM peptide samples was transferred to a 0.1 cm pathlength quartz cuvette. Scans were performed between wavelengths of 180 and 250 nm. The maximal value near 225 nm (*λ_maxHT_* _3_ = 224.2 nm) was monitored over the measured temperature range as the temperature was increased at a rate of 10 °C/hr. The Savitszky-Golay smoothing algorithm was used to calculate the derivative of the thermal melting plot. The minimum of the derivative curve describes the melting temperature (T*_m_*) of a triple helix. The molar residual ellipticity (MRE) was calculated as previously reported.^1^ All kinetics studies were performed at 0.3 mM concentration. Samples were annealed for fifteen minutes at 85 °C for fifteen minutes and then immediately measured at 5 °C for the reported period of time.

### Kinetics of HT1, HT2, and HT3

Circular dichroism was used to monitor the folding kinetics of HT1, HT2, and HT3 at 5 °C. A 3 mM peptide stock solution in 10 mM phosphate bu↵er was diluted to a working concentration of 0.3 mM peptide and 1 mM phosphate bu↵er for each sample. Samples were set to fold for a month where they were subsequently diluted to the working concentration in freshwater and then left to rest for twelve hours. A spectrum was then taken, from 210 nm to 250 nm, for each sample to yield a *λ_max_* value (see supporting information Figure S12). Each sample was then annealed at 85 °C for fifteen minutes before continuously monitoring folding for twenty-four hours at 5 °C. Samples were then monitored continuously for up to 48 hours (two days). The fraction folded for each heterotrimer that is presented in Figure 2 of the main text, is calculated by dividing the observed signal by *λ_max_* of a sample that has been folded for 700 hours (thirty days).

### Kinetics Concentration Screen of HT3

Circular dichroism was used to monitor the folding kinetics of HT3 at varying concentrations. A 3 mM peptide stock solution in 10 mM phosphate bu↵er was diluted to a working concentration. The range for e↵ective, continuous monitoring of the PPII helix (*λ ⇡* 225 nm) was determined to be from 0.4 mM to 0.15 mM. The phosphate bu↵er concentration was held consistent for each diluted sample at 1 mM. Samples were serially diluted from the stock solution and annealed at 85 °C for fifteen minutes and then immediately monitored at 5 °C for twenty-four hours. The concentration was considered in calculating the MRE for each sample. A separate sample that was folded for thirty days and then diluted to each concentration step and held for one hour to find the *λ_max_* value at the specific concentration. This value which was then used to divide and normalize the continuous folding data for the fraction folded.

#### Kinetics Temperature Screen of HT3

Circular dichroism was used to monitor the folding kinetics of HT3 at varying temperatures. A 3 mM peptide stock solution in 10 mM phosphate bu↵er was diluted to a working concentration of 0.3 mM peptide and 1mM phosphate bu↵er. A PPII helix (*λ ⇡* 225 nm) signal was monitored at temperatures in 5 °C steps, from 5 °C to 35 °C, for the peptide that was diluted to the working concentration. Each sample was annealed at 85 °C for fifteen minutes before monitoring for twenty-four hours. A separate sample that was folded for thirty days and then heated to each temperature step and held for one hour to find the *λ_max_* value at the specific temperature. This value which was then used to divide and normalize the continuous folding data for the fraction folded.

#### Comments on Fraction Folded

While the monitoring of the PPII helix formation with circular dichroism is well-reported in literature, this study of heterotrimeric folding demonstrates the need to have greater atomistic detail of collagen assembly.^2–5^ The observed signal of PPII helices can belong to varying foldons with di↵erent registers, compositions, and staggers. As a result, signals are not representative of the formation of the desired ABC-type heterotrimer. In addition, the literature reports that kinetic equations can be fit for the formation of homotrimers.^6^ However, the sequence heterogeneity of any heterotrimer makes this equation not functional. The assumption that the max signal of a trimer solution translates to a fully folded, supramolecular peptide is also not completely accurate. The conformers of each monomer, intermediate states, and other equilibrium-driven states are all contributing to the observed signal in CD. This is most obvious when comparing the observance of HT3’s formation from freely annealed monomer in Figure SS13b the raw signal is zero, but when there is a preformed heterotrimer of any other folded species the signal is falsely inflated (see Figure S9c). To prevent inaccurately measuring other states along the folding pathway, we confirmed that only the ABC-type system was observed folding along the time course with 2D HSQC NMR. Finally, for circular dichroism studies, we normalized our data to a maximum observed signal (folding signal after two weeks and equilibrating upon dilution), not as a fraction folded, for clarity of the biophysical phenomena that were observed with the kinetics studies.

#### NMR Confirms Register of ABC-Type

The register of HT3 was confirmed by using multidimensional NMR with selectively ^15^N and ^13^C isotopically labeled amino acids (see supporting information for sequence and location of isotopically labeled amino acids). Each monomer strand was labeled with a ^15^N at the Gly-21 position. Additionally, the C peptide contains both ^15^N and ^13^C label at its Gly-21, and the B peptide has the Lys-2 labeled with a ^15^N to correlate the N-terminus folding. Figure S14A shows the overlap of four di↵erent HSQC ^1^H-15N NMR spectra: one from each single peptide A, B and C, and one from the mixture of all three peptides. When combined with the data from the three 1:1 mixtures of A:B, A:C and B:C (see supporting information figure SX) this allows us to identify which peaks in the ABC mixture arise from the new assembly of three peptides (labeled with T*_x_*). These three peaks suggest that the ABC system comprises only a single composition and register. From these HSQC data, the splitting by the 13C of the most upfield signal in the 15N dimension at 106.99 ppm leads to its assignment as the C peptide. This was observed at 5 °C and 25 °C.

3D nuclear overhauser e↵ect spectroscopy (NOESY) HSQC NMR was used to determine the register of the HT3 assembly. Each assembled HT3 peak is displayed with a slice in Figure S14B and modeled in Figure S14C. The amide region of the spectra shows that the most up-field peak is the middle strand of the triple helix at 110.27 ppm in the ^15^N dimension. With the C peptide already being determined by HSQC due to splitting, the middle strand must be either the A or B peptide. The NOEs are constrained to be within 5 Å, allowing the assignment of the strands by their *↵*-protons, most importantly the hydroxyproline of the leading monomer. Using a 3D-HNHA study, the *↵*-proton of each strand’s glycine was determined (see Figure SS16). By assigning the i-1 amino acid’s *↵*-protons in Figure S14B, it is conclusive that the leading monomer can see the middle monomer’s lysine (pink circle in Figure S14B, pink line in S14c). No other correlations suggestive of different registers or compositions are observed, leading us to conclude that HT3 folds into a single composition, single register triple helix ABC in solution.

#### Curve Fitting of NMR kinetics

The 2D HSQC NMR data of Figures 4 and 5 in the main manuscript were fit with an exponential plateau function in triplicate using GraphPad Prism version 10.0.0 for Windows, GraphPad Software, Boston, Massachusetts USA, www.graphpad.com. The error was calculated on a 95% confidence interval. The constant (k) was constrained to be greater than 0. An exponential plateau function is defined with Equation 1 below.

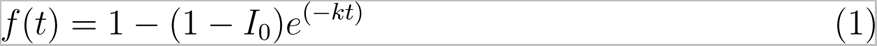

Additional analysis was carried out with the use of Bruker Dynamic Centering Software. The spectra were peak-picked for peak intensity (I) and then plots were determined to be best fit with first-order kinetics (see Equation 2 below). All confidence intervals were set to 95% as determined by weighted fit.

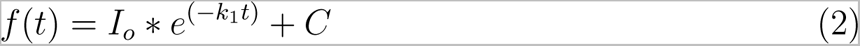

**Figure S1:**
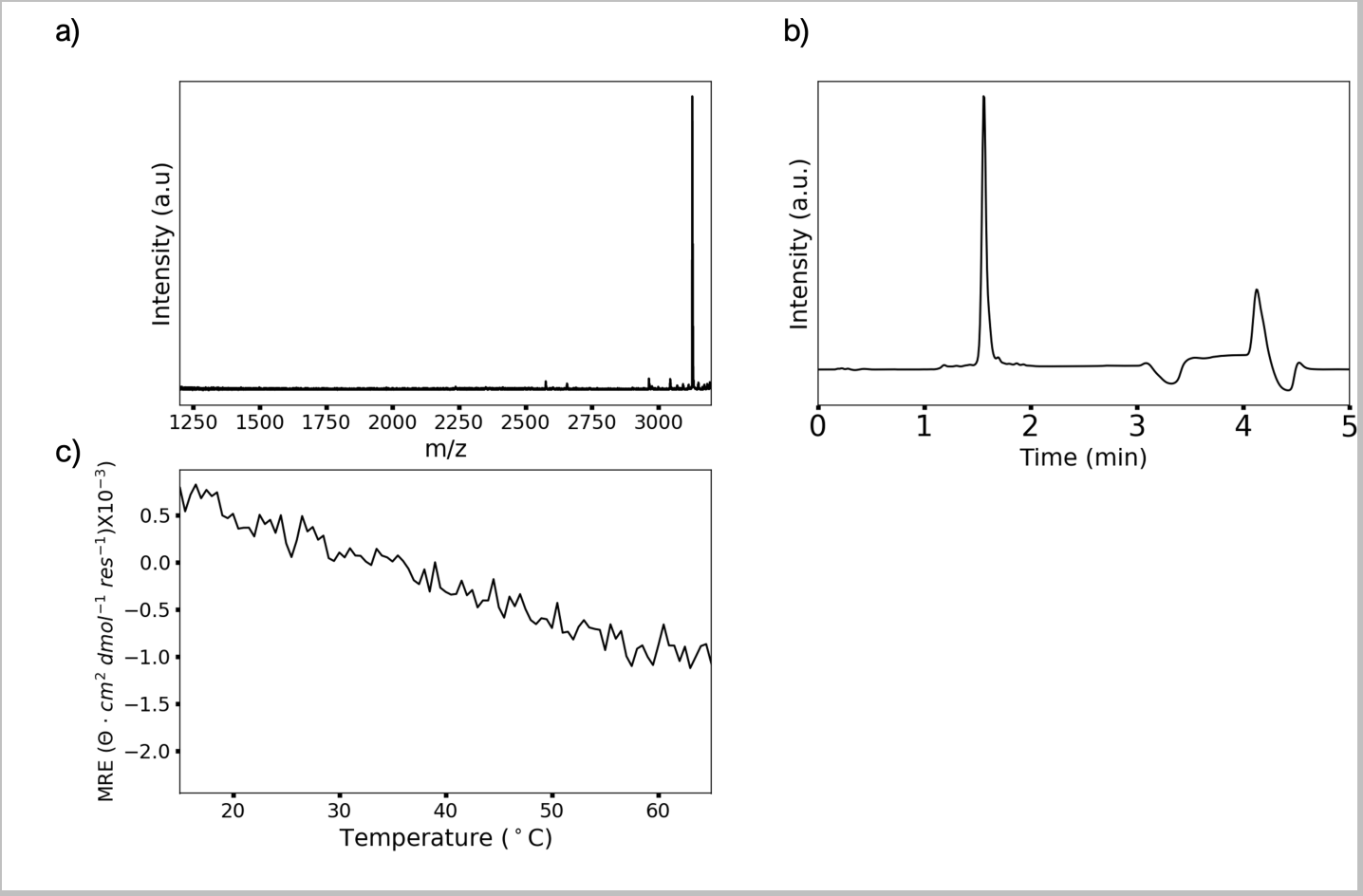
HT3A Characterization: a) mass spectra (Expected: [M + H]^+^ = 3127.67; Observed: 3127.69) b) Pure UPLC trace, CD c) melting curve

**Figure S2:**
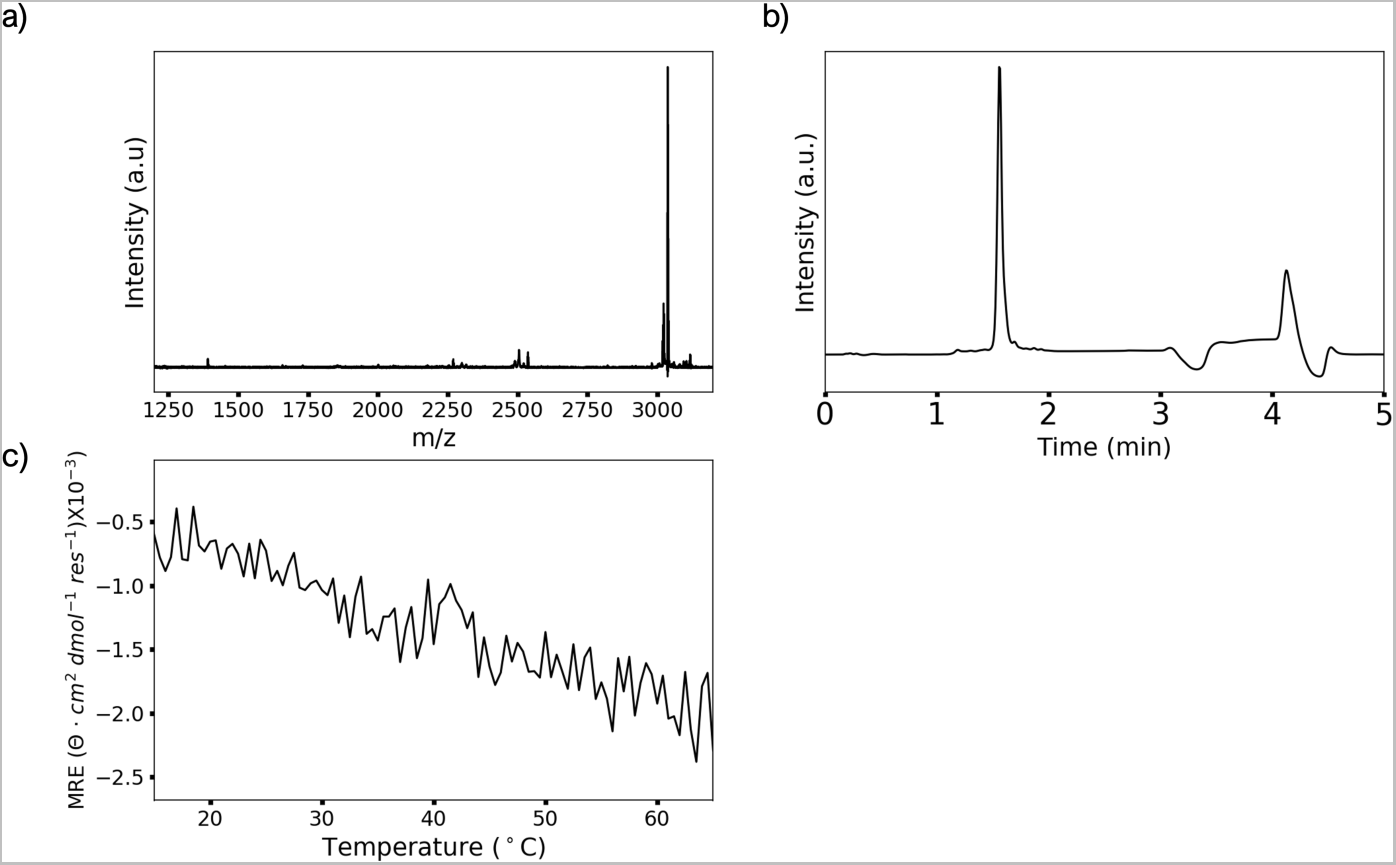
HT3B Characterization: a) mass spectra (Expected: [M + H]^+^ = 3037.48; Observed: 3037.51) b) Pure UPLC trace, CD c) melting curve

**Figure S3:**
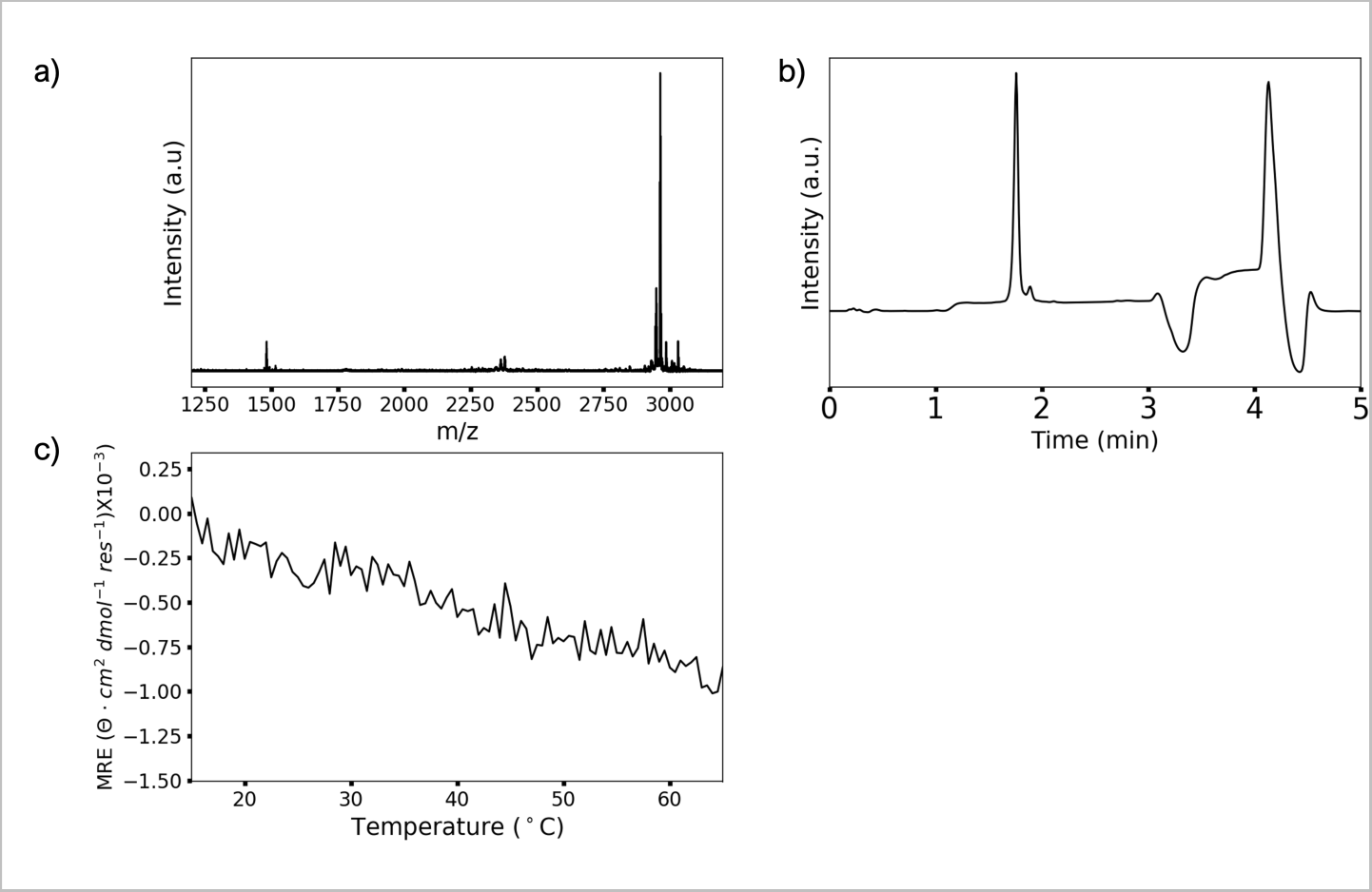
HT3C Characterization: a) mass spectra (Expected: [M + H]^+^ = 2960.24; Observed: 2962.03) b) Pure UPLC trace, CD c) melting curve

**Figure S4:**
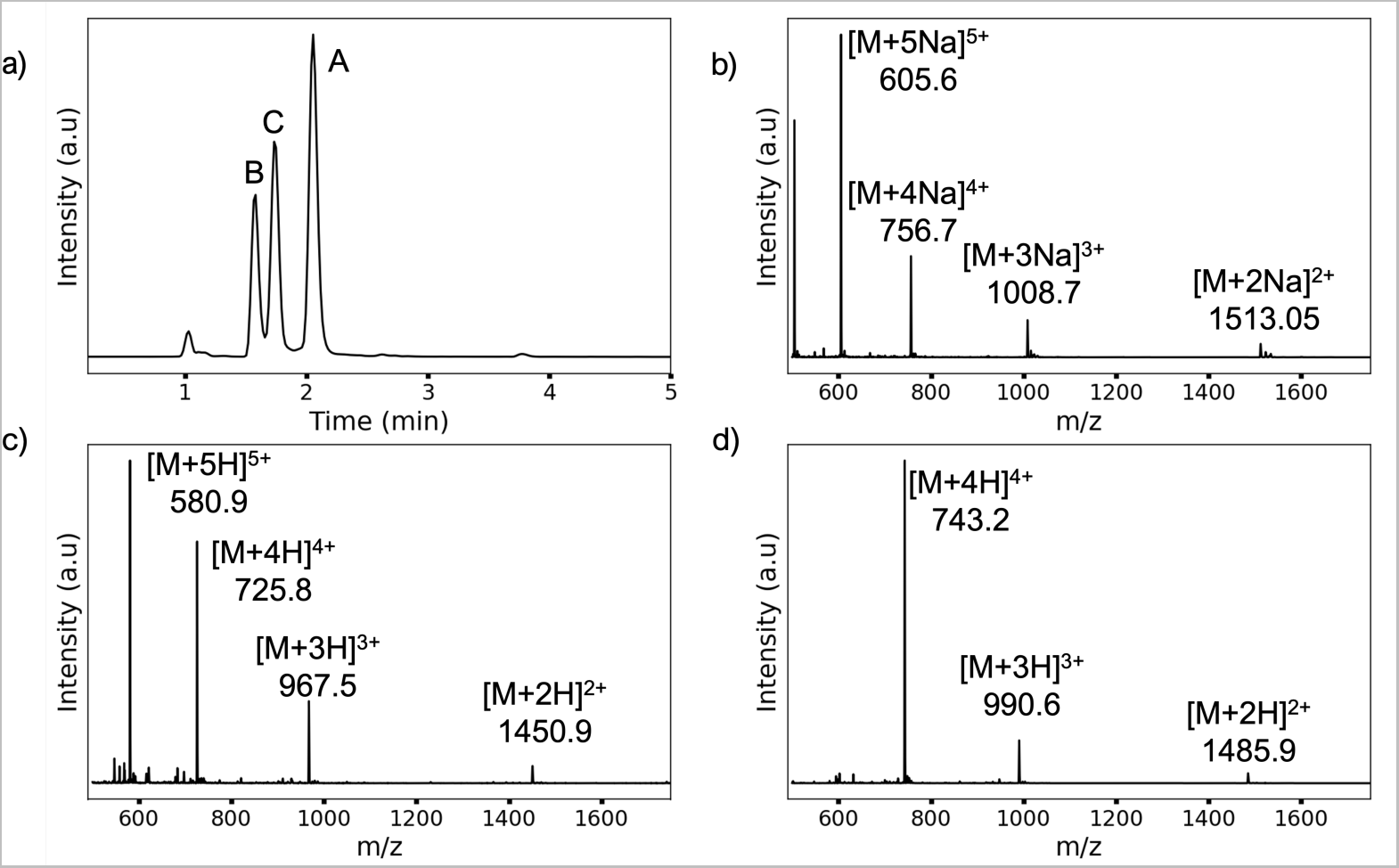
HT2 ABC LC-MS trace. a) Chromatogram of the HT2ABC Mixture. b) HT2A mass spectra: (Sequence:PKGPKGFPGPOGFKGFKGPKGPOGFKGPOG): Expected: [M + Na]^+^ = 3029; Observed: 3029.1 c) HT2B mass spectra: (Sequence:PKGDOGDKGPOGPOGDKGDOGDKGPKGDOG): Expected: [M + H]^+^ = 2901.2; Observed: 2901.8 d) b) HT2C mass spectra: (Sequence:PRGEPGPRGERGPPGPPGERGPPGEPGEPG): Expected: [M + H]^+^ = 2969.16; Observed: 2971.8

**Figure S5:**
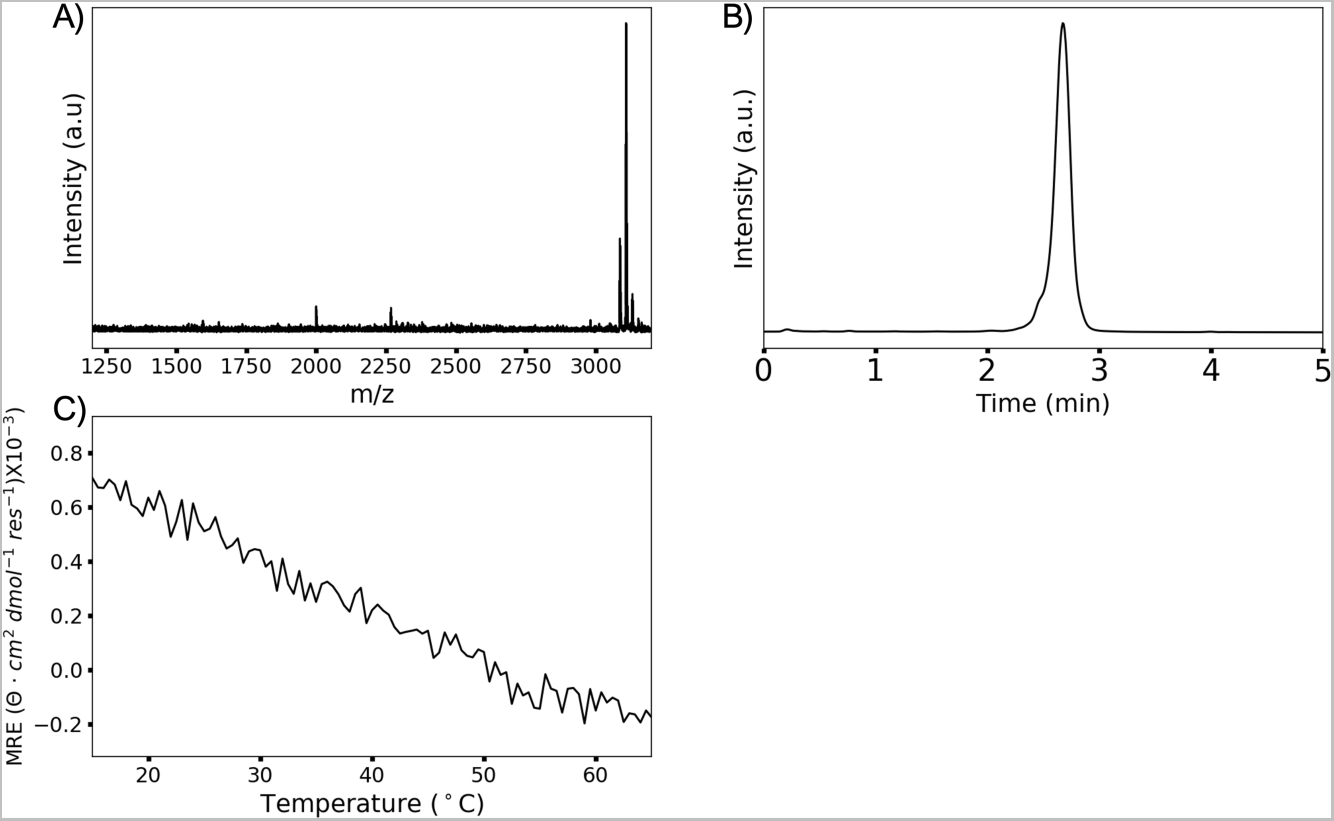
HT1A Characterization (Sequence: KPGYPGDRGPOGQRGKRGPPGDRGFPGFOG): a) mass spectra (Expected: [M + H]^+^ = 3135.58; Observed: 3136.58) b) Pure UPLC trace, CD c) melting curve

**Figure S6:**
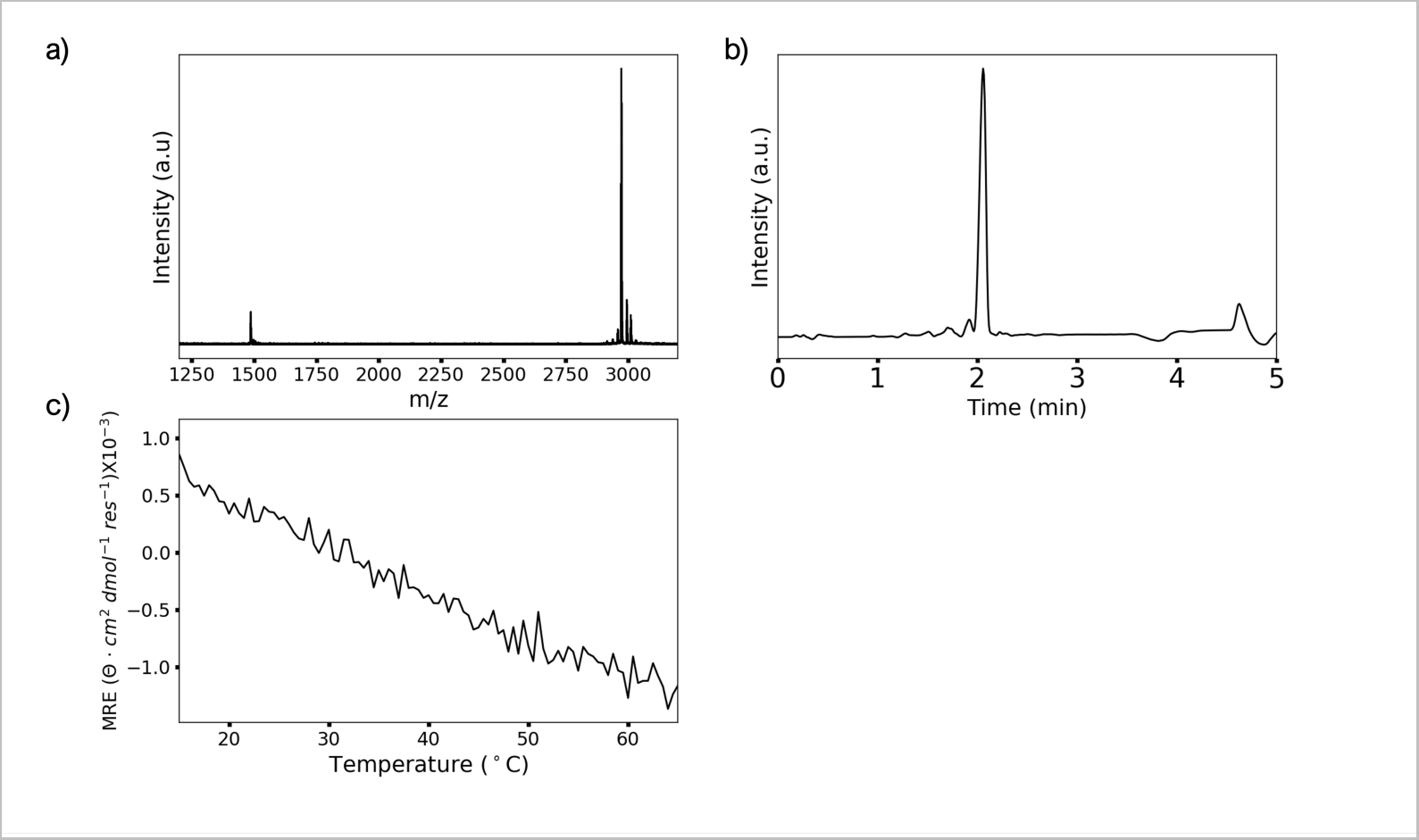
HT1B Characterization (Sequence: PRGLKGLOGAQGPOGAKGLKGPRGYK-GLOG): a) mass spectra (Expected: [M + H]^+^ = 2971.67; Observed: 2971.88) b) Pure UPLC trace, CD c) melting curve

**Figure S7:**
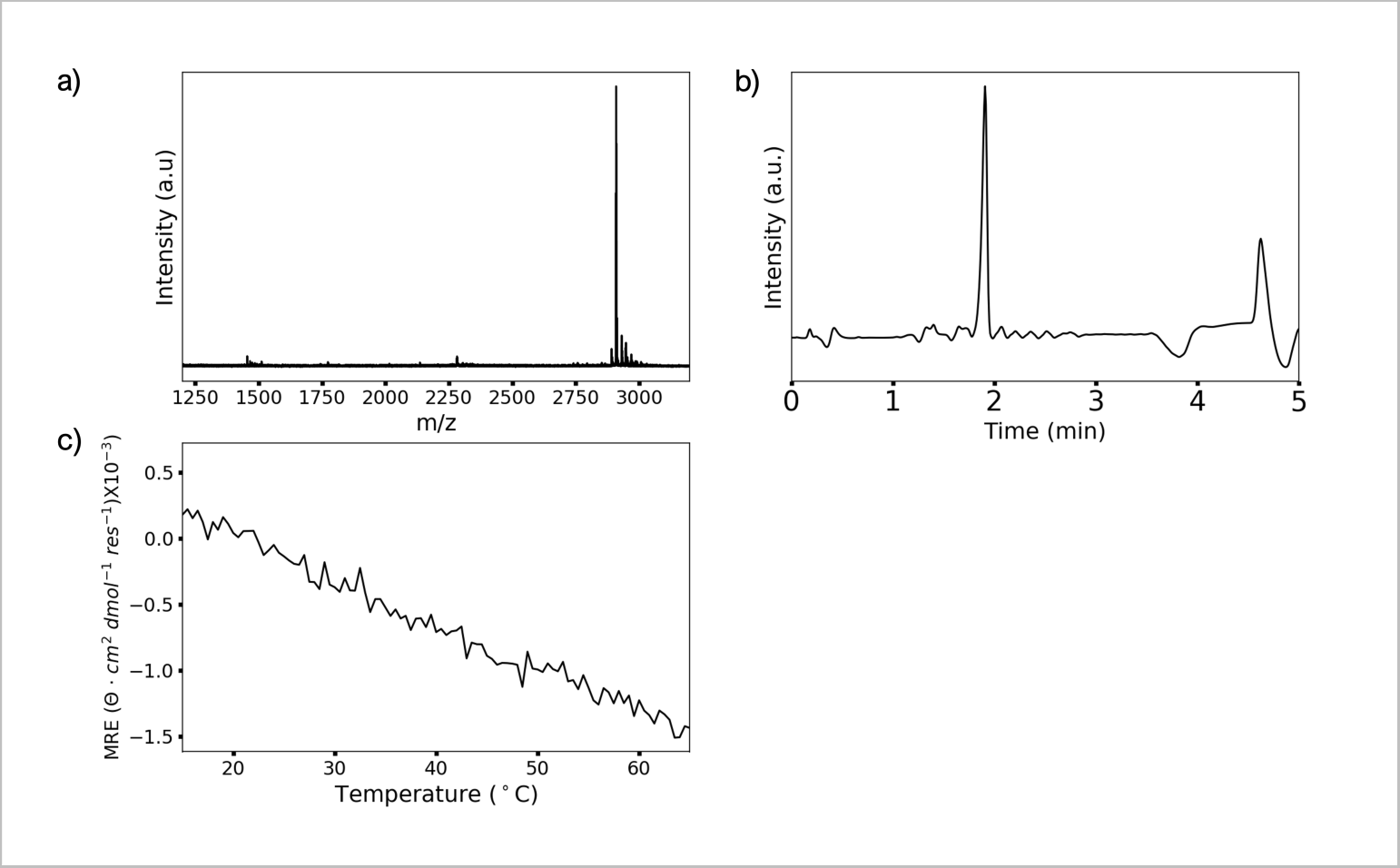
HT1C Characterization (Sequence: PKGYOGDOGIAGPDGPKGDQGDR-GAOGDOG): a) mass spectra (Expected: [M + H]^+^ = 2908.31; Observed: 2908.54) b) Pure UPLC trace, CD c) melting curve

**Figure S8:**
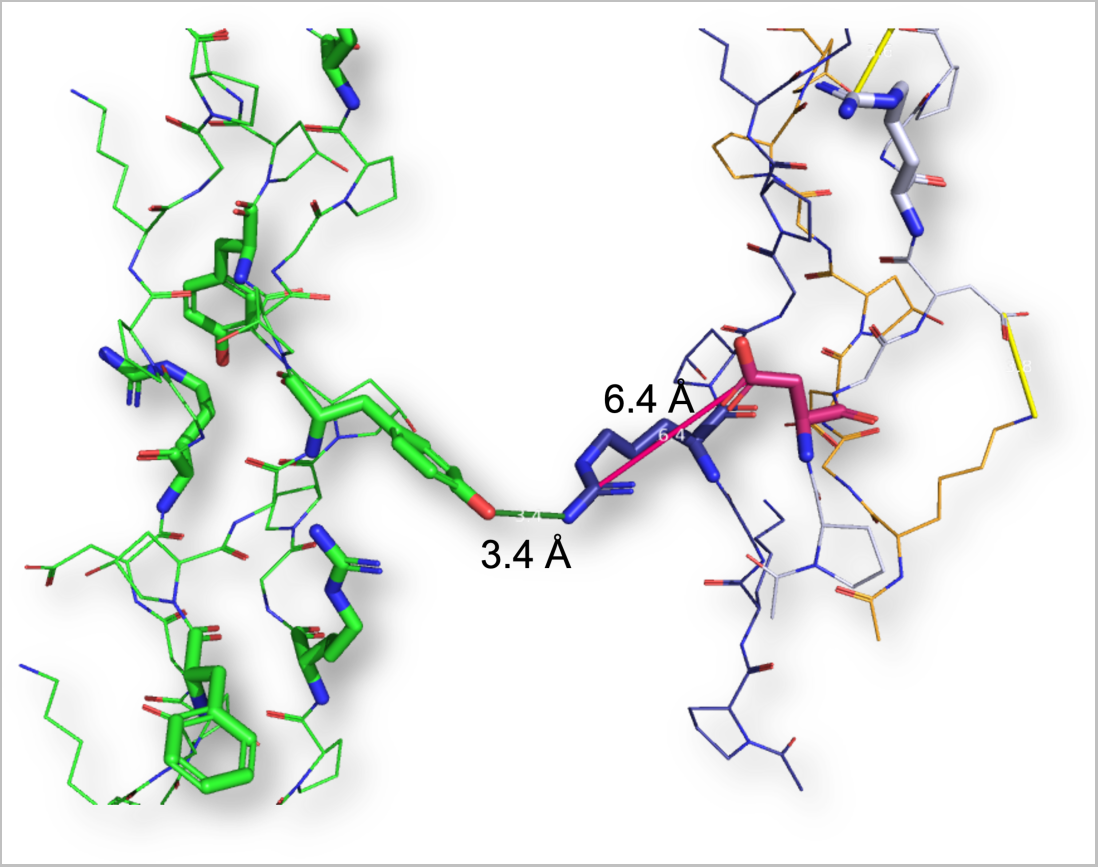
Interhelical RY Interaction. The parallel packing of the HT3 crystal structure leads to an Arg-4 (a) (Green triple helix) reaching to Tyr-13 (A’) of the blue triple helix.

**Figure S9:**
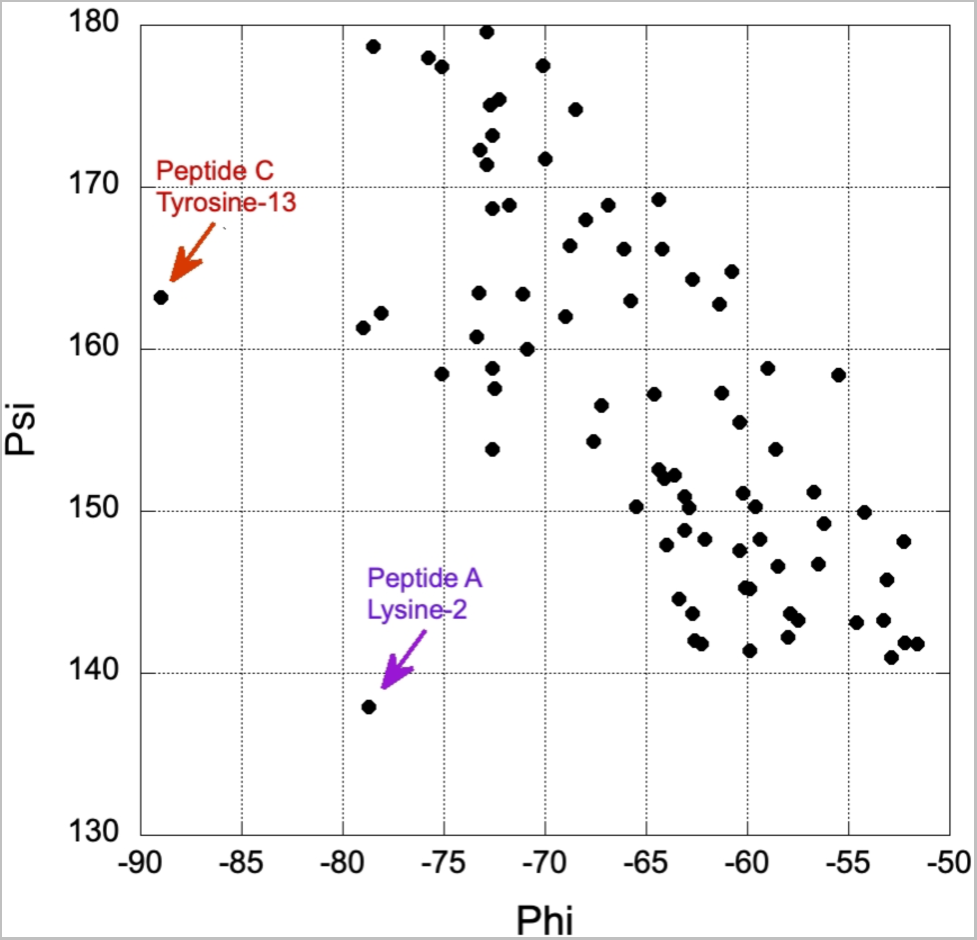
Ramachandran Plot of HT3. The phi-psi angles of each residue in the crystal structure of HT3 show good with other triple helical peptides in literature.^7^ The highest deviating residues are labeled appropriately.

**Table S1:** Pairwise Interactions in HT3.

| Interaction | Residues (Yaa-Xaa) | <sup>a</sup>Pairwise (Å) | <sup>b</sup>H-Bonding (Å) |
| --- | --- | --- | --- |
| Cation-π | Arg-11 (b) to Tyr-13 (c) | | |
| Cation-π | Arg-17 (a) to Tyr-19 (b) | 3.9 | |
| Cation-π | Arg-8 (c) to Tyr-13 (a) | 4.4 | |
| Cation-π | Arg-5 (c) to Phe-10 (a) | 3.6 | |
| Cation-π | Arg-26 (a) to Phe-28 (b) | 4.4 | |
|  | **Average Distance** | **4.1** |  |
| Lateral Charge Pair | Asp-2 (c) to Arg-4 (a) | 6.4 | 4.2 |
| Lateral Charge Pair | Asp-9 (c) to Arg-11 (a) | 4.2 | 2.9 |
| Lateral Charge Pair | Glu-20 (c) to Lys-22 (a) | 3.6 | 2.7 |
|  | **Average Distance** | **3.9** | **2.8** |
| Axial Charge Pair | Lys-2 (a) to Asp-4 (b) | 3.5 | 2.8 |
| Axial Charge Pair | Lys-8 (a) to Asp-10 (b) | 3.8 | 2.8 |
| Axial Charge Pair | Lys-23 (a) to Asp-25 (b) | 3.7 | 3.1 |
| Axial Charge Pair | Lys-2 (b) to Asp-4 (c) | 3.8 | 2.7 |
| Axial Charge Pair | Lys-14 (b) to Asp-16 (c) | 3.6 | 2.7 |
| Axial Charge Pair | Lys-20 (b) to Asp-22 (c) | 3.5 | 2.5 |
| Axial Charge Pair | Lys-26 (b) to Asp-28 (c) | 3.0 | 2.5 |
|  | **Average Distance** | **3.6** | **2.7** |

<sup>a</sup> Distance measured between the $\zeta$ -nitrogen of Lys (K) or the $\delta$ -guanidinium cation of Arg to the $\gamma$ -carbon of Asp (D) or the $\delta$ -carbon of Glu (E); <sup>b</sup> Distance measured from the $\zeta$ -nitrogen of Lys (K) or the proximal $\eta$ -Nitrogen of the guanidinium cation to the closest carboxylate oxygen of Asp (D) or Glu (E)

**Table S2:** Data collection and refinement statistics of HT3 (PDB ID 8tw0. )

| Parameter | Value |
| --- | --- |
| Space group | P 1 |
| Unit cell lengths (a,b,c, Å) | 27.70, 27.88, 28.40 |
| Unit cell angles ($\alpha, \beta, \gamma$, °) | 109.4, 111.8, 95.7 |
| Wavelength (Å) | 1.5418 |
| Temperature (K) | 100 |
| Number of sweeps / frames / degrees collected | 25 / 2508 / 1254 |
| Resolution range (Å) | 24.2 - 1.53 (1.59 - 1.53) |
| Observed reflections | 85240 (1642) |
| Unique reflections | 10535 (802) |
| Multiplicity | 8.1 (2.0) |
| Completeness (%) | 97 (74) |
| $\langle I/\sigma(I) \rangle$ | 23.4 (1.8) |
| Wilson B-factor (Å<sup>2</sup>) | 10.8 |
| $R_{merge}(I)$ | 0.080 (0.448) |
| $R_{meas}(I)$ | 0.096 (0.665) |
| $R_{pim}(I)$ | 0.029 (0.331) |
| CC<sub>1/2</sub> | 0.999 (0.714) |
| CC* | 1.00 (0.878) |
| Reflections used in refinement | 10513 (786) |
| Reflections used for R-free | 521 (35) |
| $R_{work}$ | 0.150 (0.253) |
| $R_{free}$ | 0.185 (0.263) |
| CC<sub>work</sub> | 0.979 (0.801) |
| CC<sub>free</sub> | 0.977 (0.461) |
| Number of non-hydrogen atoms | 852 |
| macromolecules | 635 |
| sulfate ions | 15 |
| water | 202 |
| Number of protein residues / chains | 89 / 3 |
| RMS<sub>bonds</sub> (Å) | 0.006 |
| RMS<sub>angles</sub> (°) | 1.07 |
| Ramachandran favored (%) | 100.0 |
| Rotamer outliers (%) | 0.0 |
| Clashscore | 0.8 |
| Average B-factor (Å<sup>2</sup>) | 17.7 |
| macromolecules (Å<sup>2</sup>) | 15.5 |
| sulfate ions (Å<sup>2</sup>) | 28.2 |
| water (Å<sup>2</sup>) | 23.8 |
| Number of TLS groups | 11 |
| *Statistics for the highest-resolution shell are shown in parentheses.* |  |

**Figure S10:**
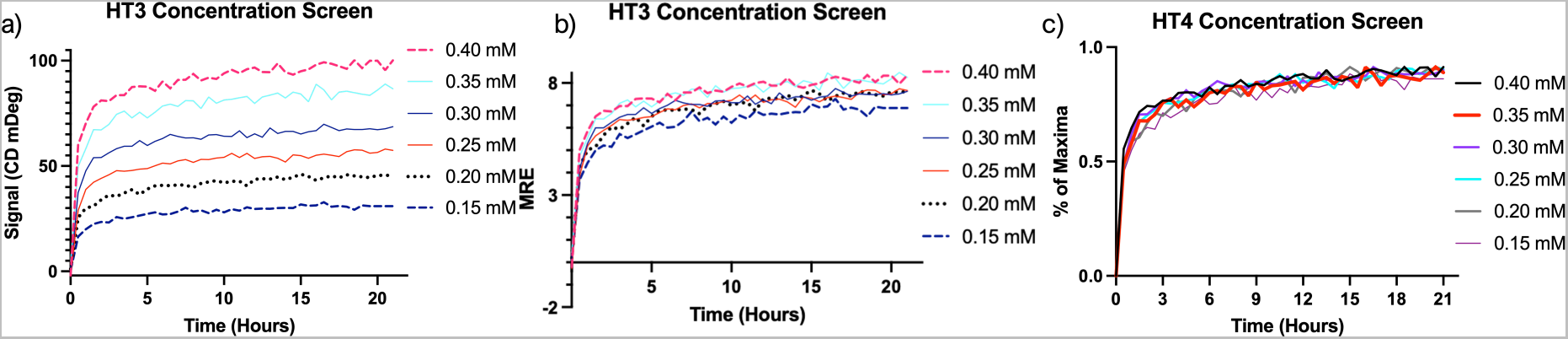
Circular dichroism of HT3 at Varying Concentrations. a) Non-converted signal of HT3 formation for varying concentrations. b) Converted MRE values at varying concentrations c) Fraction of maxima that is normalized with the maximum signal value for each sample’s concentration.

**Figure S11:**
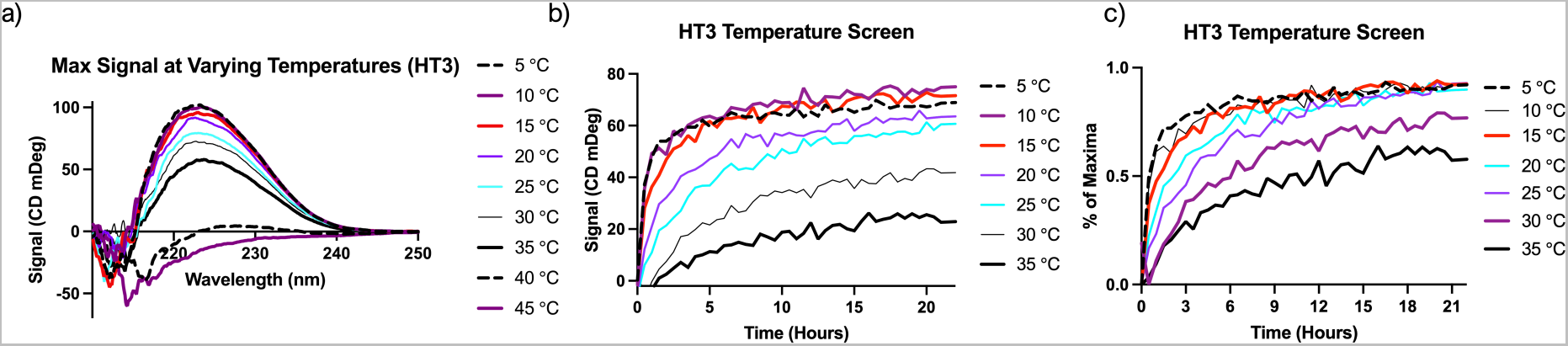
Circular dichroism of HT3 at Varying Temperatures. a) A CD scan of HT3 at varying temperatures indicates di↵erent maxima depending on the temperature b) Non-converted signal of HT3 formation for varying concentrations. c) Non-converted signal of HT3 formation for temperature scans

**Figure S12:**
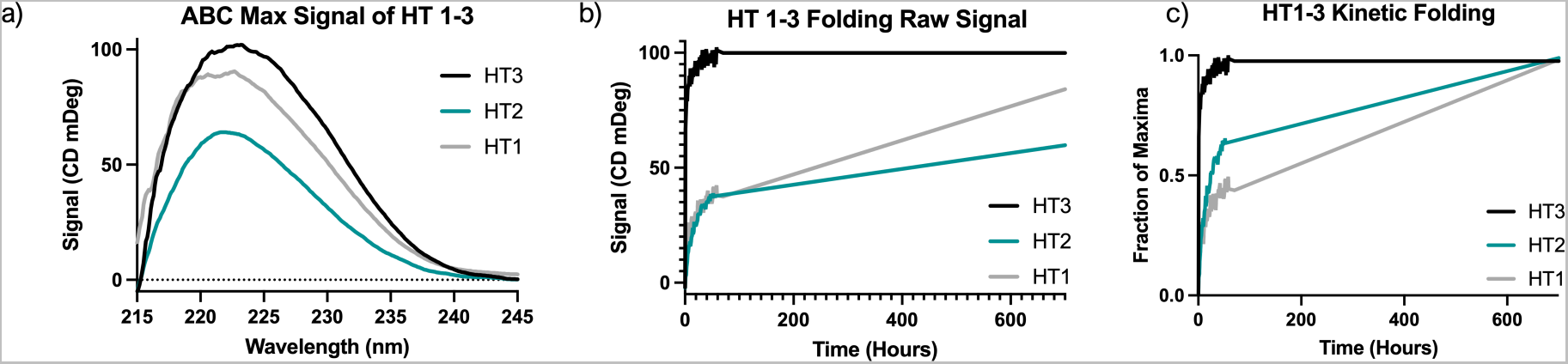
Circular dichroism of ABC Mixtures of HT1, HT2, and HT3 at Long Time Scales. a) A CD scan of HT1-3 at day 30 indicates di↵erent maxima depending on the heterotrimer b) Non-converted signal of HT1-3 formation. c) Converted signal of HT1-3 where the kinetic folds are divided by the maxima of each ABC heterotrimer after 30 days of folding.

**Figure S13:**
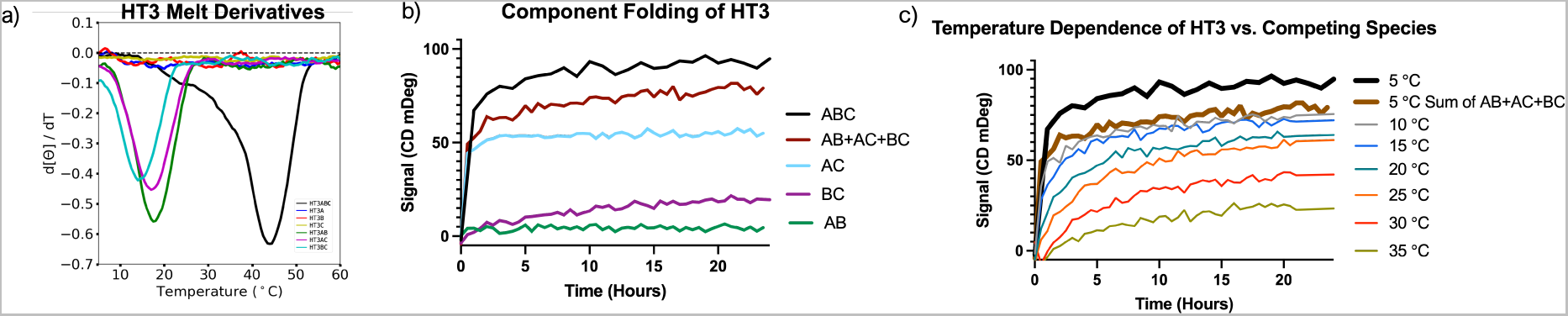
Circular dichroism of HT3 Compositions. a) A CD melt curve derivative scan of HT3 and its varying mixtures that comprise the ABC heterotrimer b) Non-converted signal of the binary component mixtures and their summed values (AB+AC+Bc) compared to the folding of HT3 ABC at 5 °C. c) A comparison of ABC folding at various temperatures, from 25-35 °C other species are unfolded, but there is still a temperature dependence.

**Table S3:** Circular Dichroism Kinetics of HT3 at Varying Concentrations.

| Concentration (mM) | Percent Folded | Time (Hours) |
| --- | --- | --- |
| 0.15 mM | 50% | 0.53 |
|  | 75% | 2.83 |
| 0.20 mM | 50% | 0.53 |
|  | 75% | 2.28 |
| 0.25 mM | 50% | 0.53 |
|  | 75% | 2.34 |
| 0.30 mM | 50% | 0.50 |
|  | 75% | 2.12 |
| 0.35 mM | 50% | 0.53 |
|  | 75% | 2.29 |
| 0.40 mM | 50% | 0.37 |
|  | 75% | 1.65 |

<sup>a</sup> All experimental values were determined at 5 °C

**Table S4:** Circular Dichroism Statistics of HT3 at Varying Concentrations.

|  | 0.40 mM | 0.35 mM | 0.30 mM | 0.25 mM | 0.20 mM | 0.15 mM |
| --- | --- | --- | --- | --- | --- | --- |
| k (hr<sup>-1</sup>) | 1.358 | 0.9855 | 1.124 | 1.008 | 0.6788 | 0.6890 |
| **95% CI** |  |  |  |  |  |  |
| k (hr<sup>-1</sup>) | 0.99 – 1.85 | 0.72 – 1.34 | 0.88 – 1.42 | 0.76 – 1.33 | 0.47 – 0.99 | 0.49 – 0.98 |
| R<sup>2</sup> | 0.9063 | 0.9061 | 0.9340 | 0.9185 | 0.8875 | 0.8948 |

<sup>a</sup> All experimental values were determined at 5 °C and fit with a second exponential equation and at a 95% confidence interval. Samples were performed at n = 1.

**Table S5:** Circular Dichroism Kinetics of HT3 at Varying Temperatures.

| Temperature (°C) | Percent Folded | Time (Hours) | $\lambda_{max}$ (CD mDeg) |
| --- | --- | --- | --- |
| 5 °C | 50% | 0.50 |  |
|  | 75% | 2.12 | 99.5 |
| 10 °C | 50% | 0.50 |  |
|  | 75% | 3.68 | 99.9 |
| 15 °C | 50% | 1.25 |  |
|  | 75% | 4.35 | 94.3 |
| 20 °C | 50% | 2.30 |  |
|  | 75% | 6.21 | 89.1 |
| 25 °C | 50% | 3.37 |  |
|  | 75% | 8.85 | 78.6 |
| 30 °C | 50% | 6.15 |  |
|  | 75% | 17.00 | 70.9 |
| 35 °C | 50% | 11.92 | 57.5 |

<sup>a</sup>All experimental values were determined at 0.30 mM

**Table S6:** Circular Dichroism Statistics of HT3 at Varying Temperatures.

|  | 5 °C | 10 °C | 15 °C | 20 °C | 25 °C | 30 °C | 35 °C |
| --- | --- | --- | --- | --- | --- | --- | --- |
| k (hr<sup>-1</sup>) | 1.124 | 0.5799 | 0.4167 | 0.2859 | 0.1935 | 0.1324 | 0.07129 |
| **95% CI** |  |  |  |  |  |  |  |
| k (hr<sup>-1</sup>) | 0.88–1.42 | 0.40–0.85 | 0.33–0.52 | 0.24–0.34 | 0.17–0.22 | 0.10–0.17 | 0.05–0.10 |
| R<sup>2</sup> | 0.9340 | 0.8754 | 0.9325 | 0.9575 | 0.9783 | 0.9491 | 0.9193 |

<sup>a</sup> All experimental values were determined at 0.3 mM and fit with a second exponential equation and at a 95% confidence interval. Samples were performed at n = 1.

**Table S7:** Kinetics of HT3-ABC Formation with NMR.

| Sample | A (k<sub>1</sub> min<sup>-1</sup>) | B (k<sub>1</sub> min<sup>-1</sup>) | C (k<sub>1</sub> min<sup>-1</sup>) | N-term (k<sub>1</sub> min<sup>-1</sup>) |
| --- | --- | --- | --- | --- |
| A+B+C - 1 | 0.0125 ± 0.0012 | 0.0071 ± 0.0012 | 0.0054 ± 0.0012 | 0.0082 ± 0.0010 |
| A+B+C - 2 | 0.0085 ± 0.0012 | 0.0097 ± 0.0016 | 0.0081 ± 0.0013 | 0.0082 ± 0.0011 |
| A+B+C - 3 | 0.0360 ± 0.0064 | 0.0496 ± 0.0044 | 0.0529 ± 0.0042 | 0.0468 ± 0.0091 |
| **Average** | **0.0190 ± 0.0030** | **0.0212 ± 0.0024** | **0.0210 ± 0.0037** | **0.0211 ± 0.0037** |
| BC+A - 1 | 0.0045 ± 0.0014 | 0.0048 ± 0.0016 | 0.0045 ± 0.0019 | 0.0047 ± 0.0012 |
| BC+A - 2 | 0.0039 ± 0.0008 | 0.0042 ± 0.0010 | 0.0044 ± 0.0009 | 0.0043 ± 0.0007 |
| BC+A - 3 | 0.0067 ± 0.0008 | 0.0055 ± 0.0007 | 0.0060 ± 0.0007 | 0.0069 ± 0.0007 |
| **Average** | **0.0050 ± 0.0010** | **0.0049 ± 0.0011** | **0.0050 ± 0.0011** | **0.0053 ± 0.0009** |
| AB+C | 0.0491 ± 0.0145 | 0.0612 ± 0.0265 | 0.0590 ± 0.0177 | 0.0146 ± 0.0094 |
| AC+B | 0.0092 ± 0.0030 | 0.0876 ± 0.0520 | 0.0027 ± 0.0021 | 0.0039 ± 0.0021 |

<sup>a</sup> All experimental values were determined at 2.8 mM peptide diluted in 10% (v/v) D<sub>2</sub>O in 10 mM phosphate buffer at 5 °C

**Table S8:** Kinetics of HT3-ABC Formation with NMR at Various Temperatures.

| Sample | A (k<sub>1</sub> min<sup>-1</sup>) | B (k<sub>1</sub> min<sup>-1</sup>) | C (k<sub>1</sub> min<sup>-1</sup>) | N-term (k<sub>1</sub> min<sup>-1</sup>) |
| --- | --- | --- | --- | --- |
| A+B+C-5 °C<sup>b</sup> | **0.0190 ± 0.0030** | **0.0212 ± 0.0024** | **0.0210 ± 0.0037** | **0.0211 ± 0.0037** |
| A+B+C-15°C | 0.0173 ± 0.0004 | 0.0158 ± 0.0004 | 0.0150 ± 0.0004 | 0.0141 ± 0.0003 |
| A+B+C-25 °C | 0.0087 ± 0.0001 | 0.0104 ± 0.0002 | 0.0024 ± 0.0003 | 0.0086 ± 0.0008 |
| A+B+C-30°C-1 | 0.0073 ± 0.0001 | 0.0077 ± 0.0002 | 0.0072 ± 0.0001 | 0.0069 ± 0.0003 |
| A+B+C-30°C-2 | 0.0079 ± 0.0002 | 0.0077 ± 0.0001 | 0.0072 ± 0.0001 | 0.0075 ± 0.0002 |
| A+B+C-30°C-3 | 0.0079 ± 0.0003 | 0.0069 ± 0.0002 | 0.0077 ± 0.0002 | 0.0062 ± 0.0005 |
| **30°C Average<sup>b</sup>** | **0.0077 ± 0.0002** | **0.0074 ± 0.0002** | **0.0074 ± 0.0001** | **0.0072 ± 0.0003** |

<sup>a</sup> All experimental values were determined at 2.8 mM peptide diluted in 10% (v/v) D<sub>2</sub>O in 10 mM phosphate buffer; <sup>b</sup> These are average folding k-values of triplicate independent trials.

**Figure S14:**
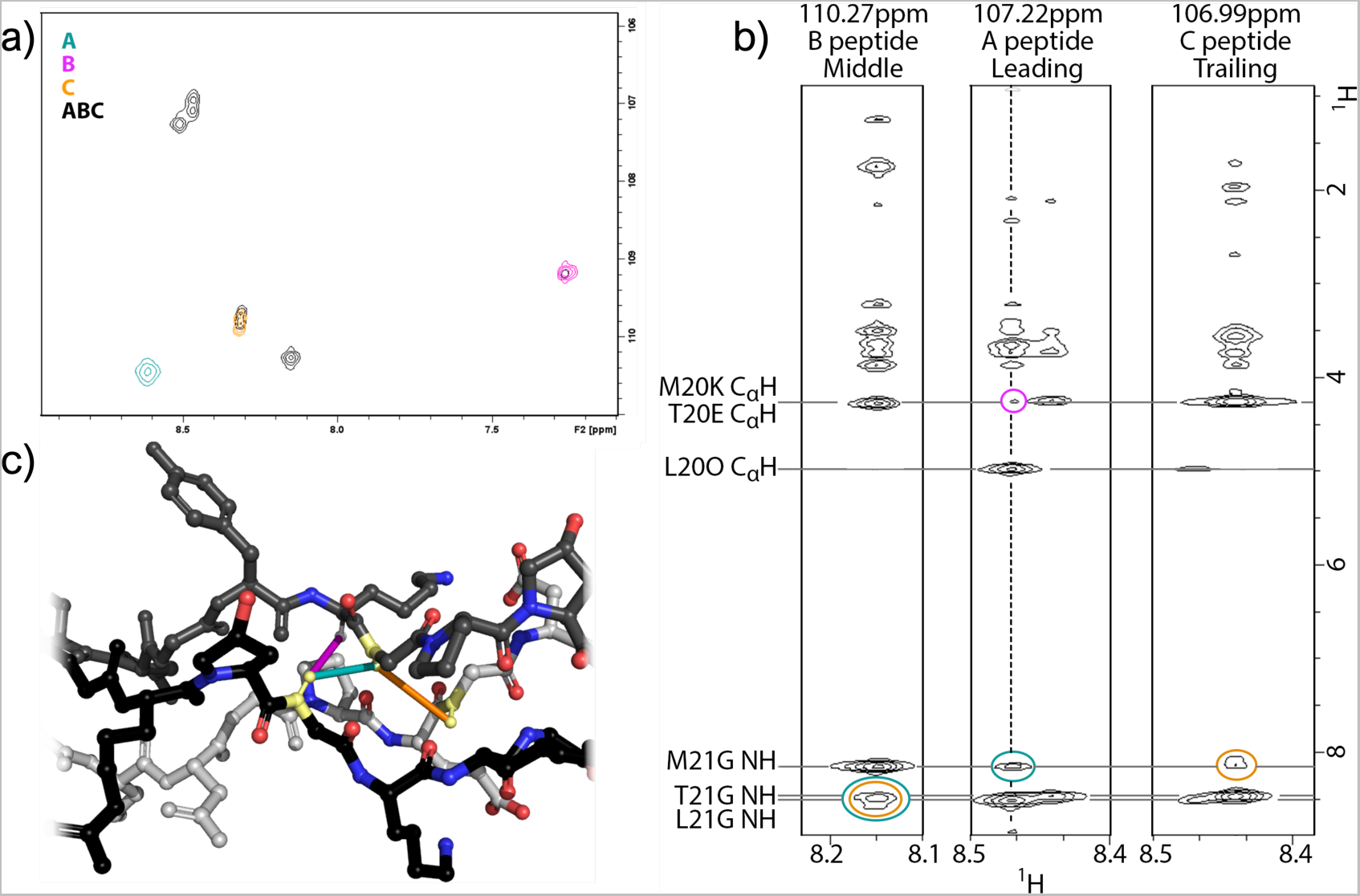
Confirmation of the ABC Register in solution for HT3 a) Heteronuclear single quantum coherence spectroscopy (HSQc) NMR of unary mixtures overlaid with the ABC-type heterotrimer mixture b) NOESY-HSQC slices of each heterotrimer peak determines an ABC register c) A 3D rendering of the NOE’s of each proton correlated to their corresponding peaks. The leading, middle, and trailing strands are black, gray, and white, respectively. Spectra were obtained at 25°C.

**Figure S15:**
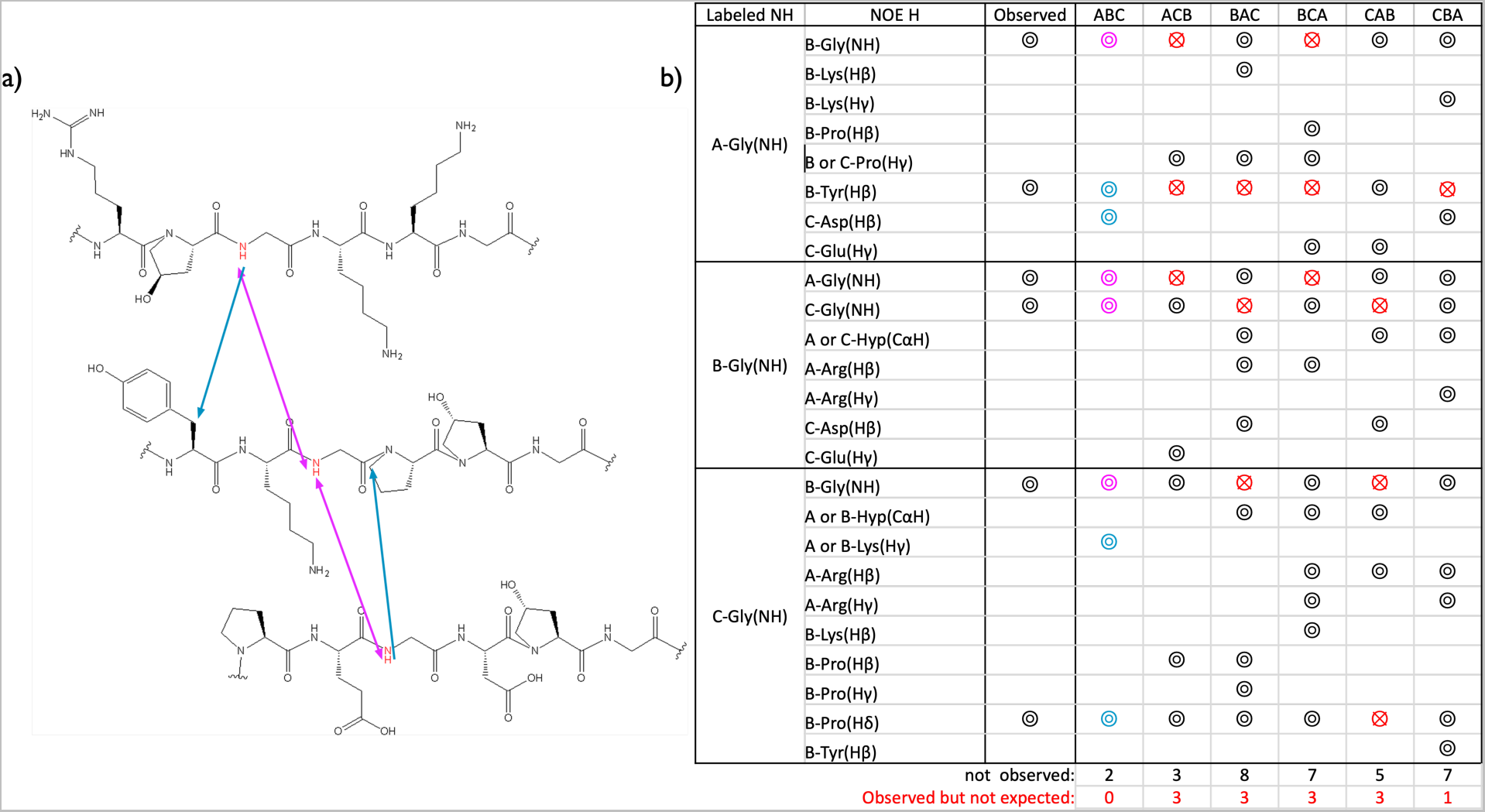
Register NOE overlap for HT3. a) A schematic of the NOE region in the leading, middle, and trailing strands of HT3. Arrows indicating the observed NOE signal as observed in the 3D NOESY-HSQC experiment. b) A tabulated presentation of di↵erent registers with their respective NOEs correlated to the spectra in Figure SS14.

**Figure S16:**
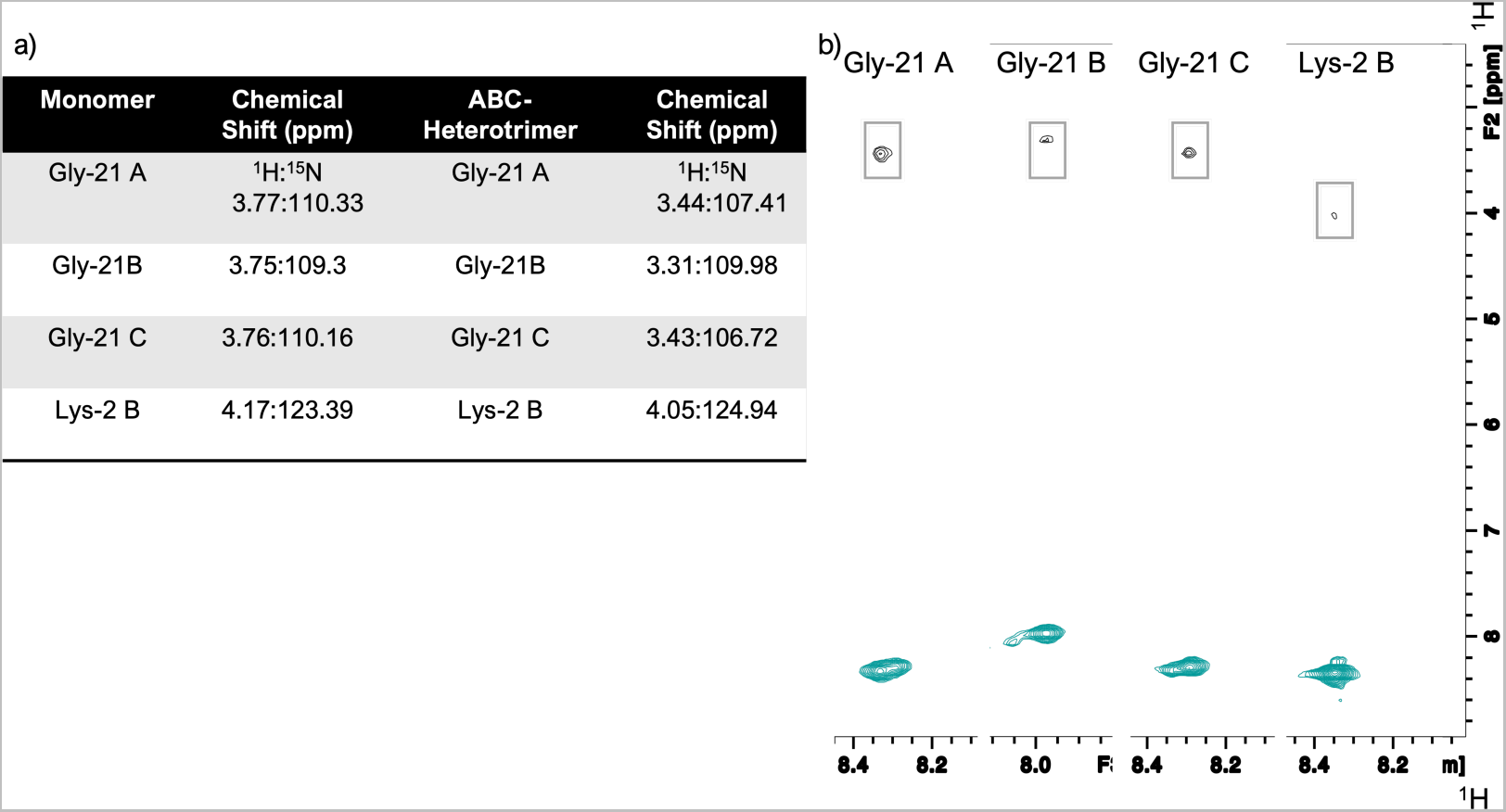
3D HNHA Confirms Proton Shifts a) A table summarizing the chemical shifts for each *↵*-proton of the labeled strands b) Slices of each correlative HNHA signal all of the labeled amino acids in the ABC-heterotrimers. Spectra were obtained at 5°C.

**Figure S17:**
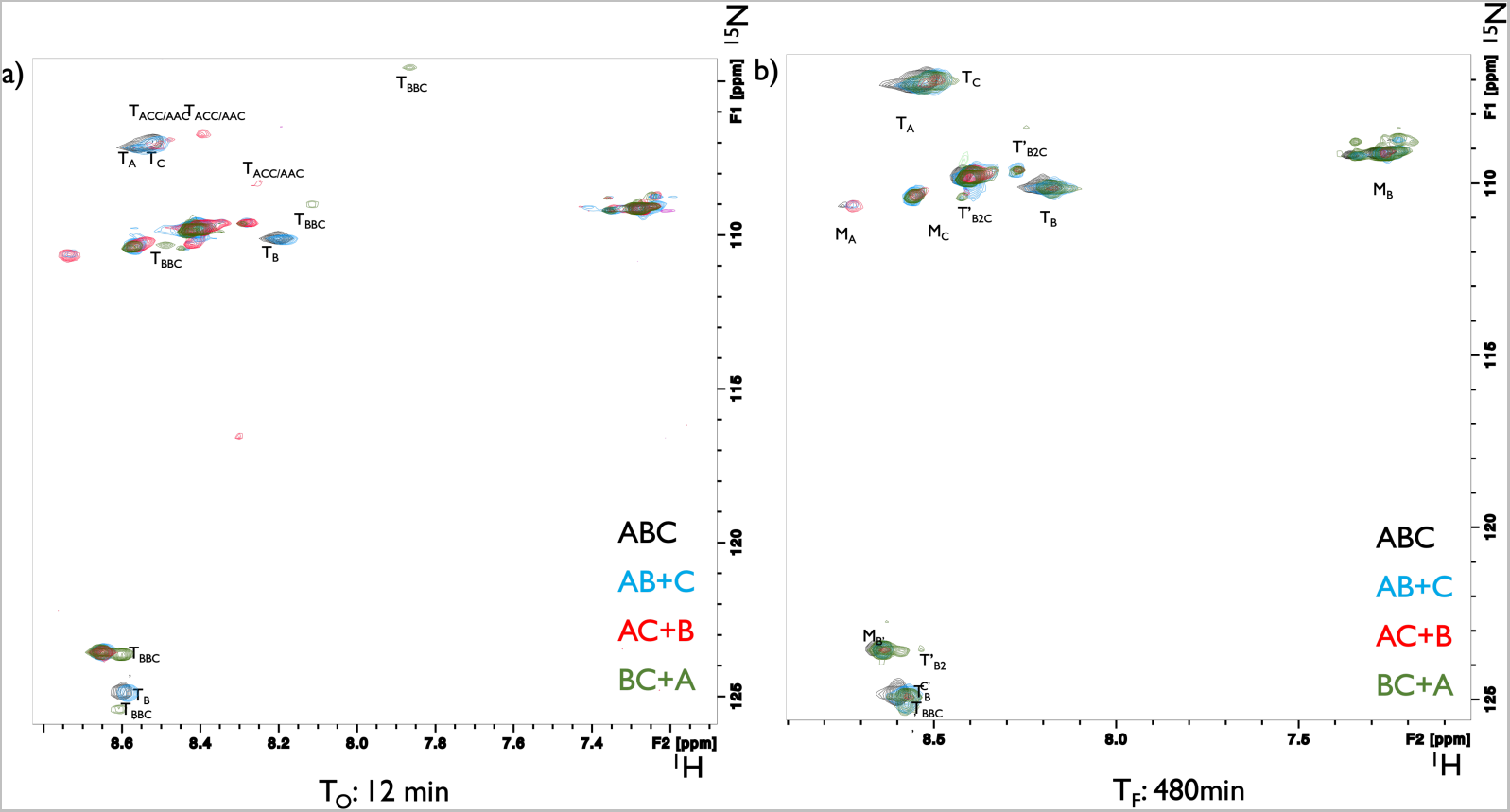
Binary mixture folding experiments via HSQC-NMR. a) The T_0_ (12 min) HSQC spectra of each mixture with an additional component that folds into HT3ABC. Two peaks from the B-strand’s 2-Lys labeling suggest a B_2_C composition b) The T*_f_* (400 min) for each mixture with an additional component that folds into HT3ABC. Spectra were obtained at 5°C.

**Figure S18:**
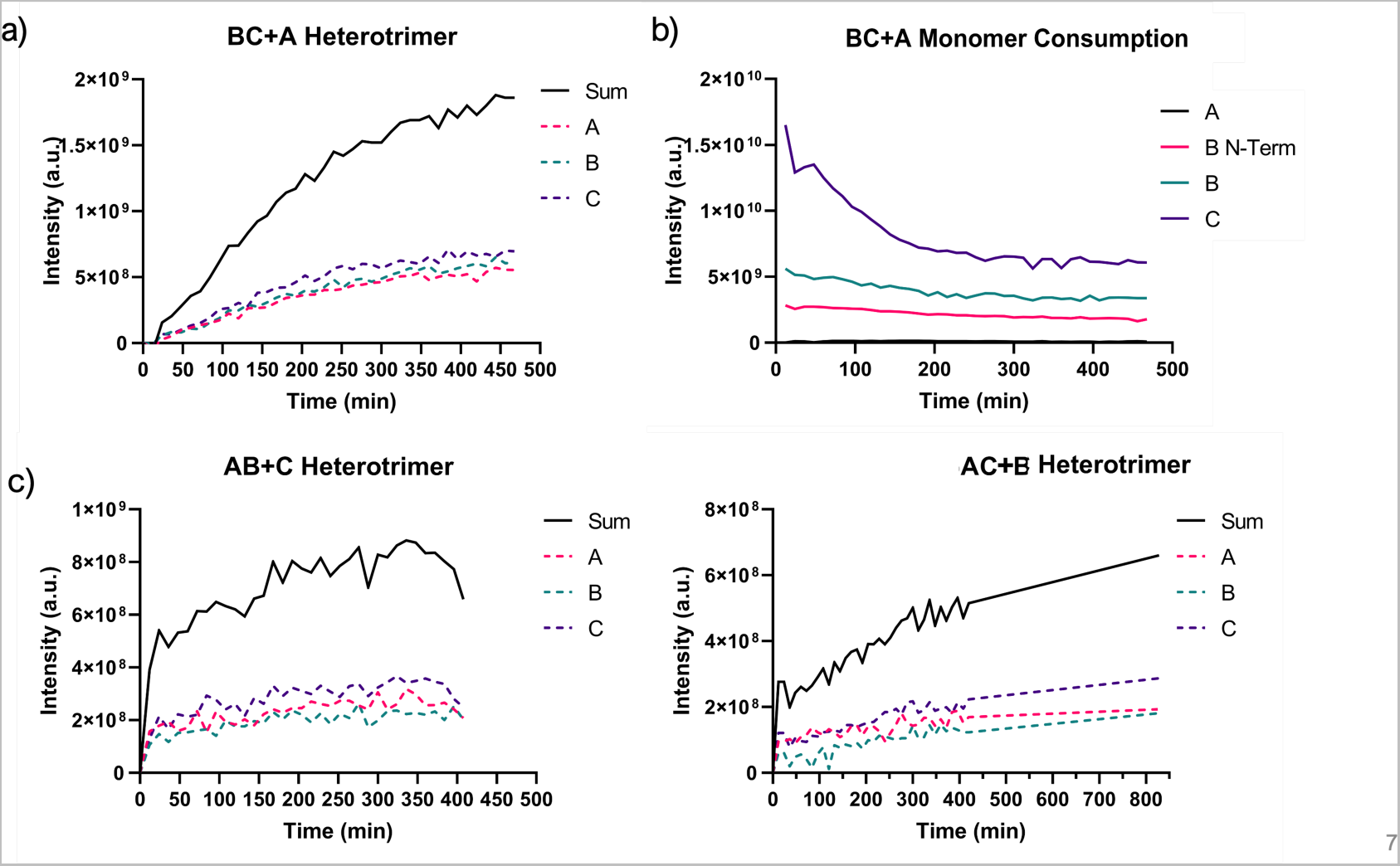
Binary mixture folding plots. a) The formation of HT3ABC from a preformed BC mixture that has monomer A added to it b) Consumption of each monomer in solution c) The formation of HT3ABC from a preformed mixture of AB heterotrimer with the addition of a free C monomer d) The formation of HT3ABC from a preformed AC heterotrimer with the addition of the free B monomer. Spectra were obtained at 5°C.

**Figure S19:**
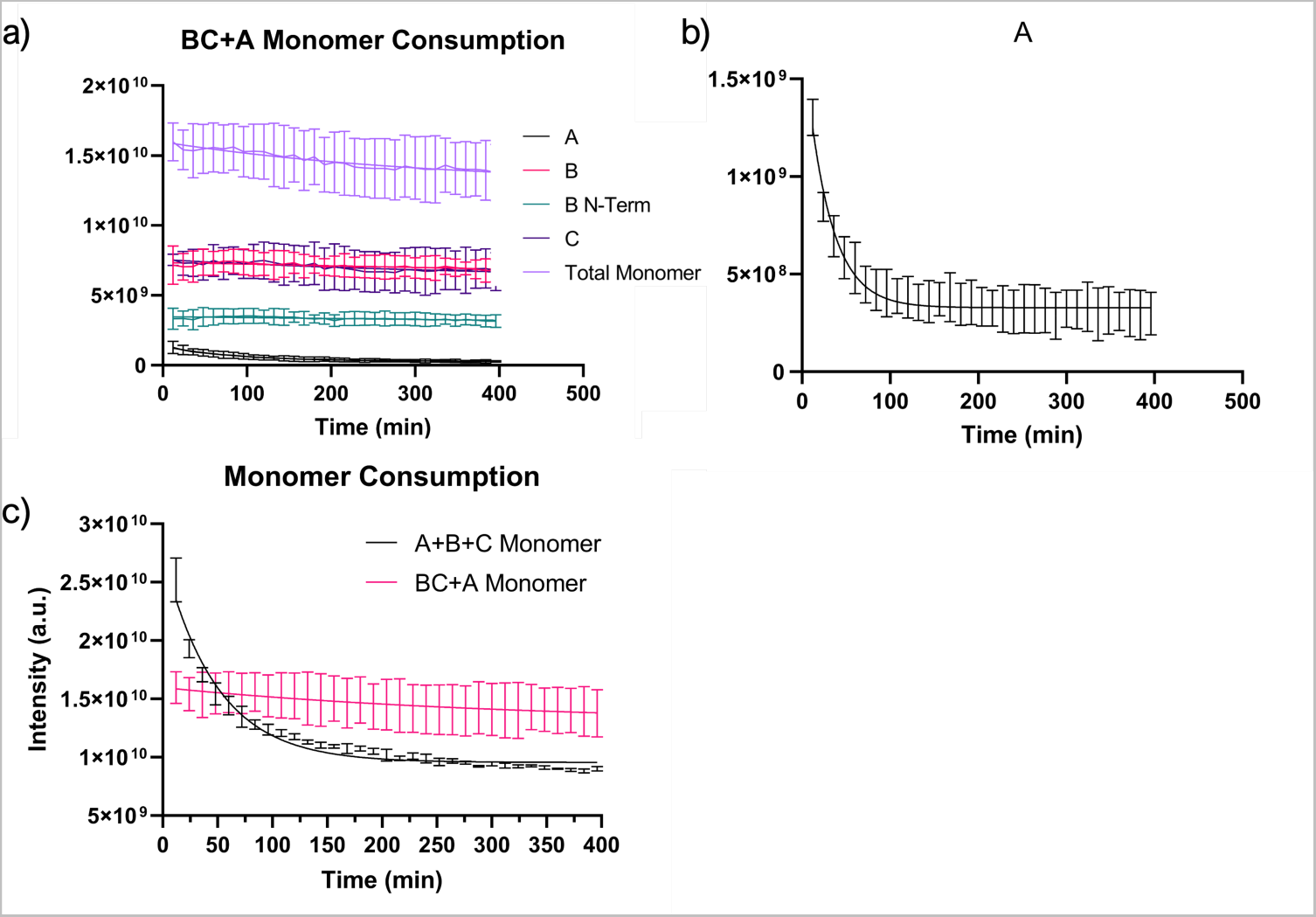
Monomer consumption when folding HT3ABC. a) The varying decrease in monomer signals in a preformed BC mixture with an addition of A monomer b) The consumption of A monomer in the BC+A mixture c) Comparison of the total monomer consumption in a free-forming mixture (A+B+c) to a preformed BC heterotrimer (BC+a). Spectra were obtained at 5°C.

**Figure S20:**
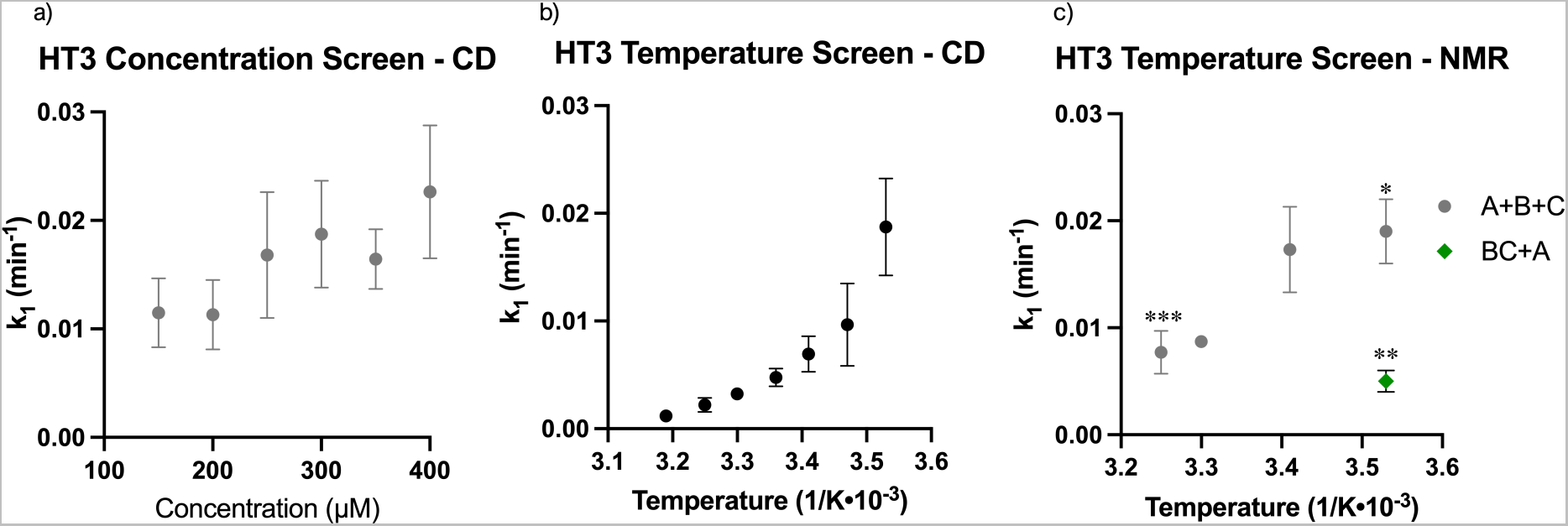
Plots of k values from Circular Dichroism and HSQC-NMR. a) A plot of k from fitting a non-linear exponential function (equation 1) of the folding curves at various concentrations of HT3 in the main Figure 2f and Tables SS3 and SS4 b) A plot of k from fitting a non-linear exponential function (equation 1) of the folding curves of the temperature screen by CD in the main Figure 2g and Tables SS5 and SS6 c) A plot of k_1_ from fitting a first order function (equation 2) of the folding curves in the main Figure 4d and Tables SS7 and SS8. *,**, and *** Indicates a sample that was done in triplicate and is noted as significantly di↵erent (on a 95% C.I.) than the other two samples that were also examined in triplicate.

## Notes

### Competing Interest Statement

The authors have declared no competing interest.

## References

(1) Lukomski, S.; Bachert, B. A.; Squeglia, F.; Berisio, R. Collagen-like proteins of pathogenic streptococci. Mol. Microbiol. 2017, 103, 919–930.

(2) Thomas, A. H.; Edelman, E. R.; Stultz, C. M. Collagen fragments modulate innate immunity. *Experimental Biology and Medicine (Maywood*, N.J.) 2007, 232, 406–411.

(3) Gordon, M. K.; Hahn, R. A. Collagens. Cell Tissue Res. 2009, 339, 247.

(4) Fratzl, P.; Weinkamer, R. Nature’s hierarchical materials. Progress in Materials Science 2007, 52, 1263–1334.

(5) Abendstein, L.; Dijkstra, D. J.; Tjokrodirijo, R. T.; van Veelen, P. A.; Trouw, L. A.; Hensbergen, P. J.; Sharp, T. H. Complement is activated by elevated IgG3 hexameric platforms and deposits C4b onto distinct antibody domains. Nat. Comm. 2023, 14, 4027.

(6) Yu, L. T.; Hancu, M. C.; Kreutzberger, M. A.; Henrickson, A.; Demeler, B.; Egelman, E. H.; Hartgerink, J. D. Hollow Octadecameric Self-Assembly of Collagen-like Peptides. J. Am. Chem. Soc. 2023, 145, 5285–5296.

(7) Shoulders, M. D.; Raines, R. T. Collagen Structure and Stability. Annu. Rev. Biochem. 2009, 78, 929–958, Publisher: Annual Reviews.

(8) Brodsky, B.; Persikov, A. V. Advances in Protein Chemistry; Fibrous Proteins: Coiled-Coils, Collagen and Elastomers; Academic Press, 2005; Vol. 70; pp 301–339.

(9) Shoulders, M. D.; Guzei, I. A.; Raines, R. T. 4-Chloroprolines: Synthesis, conformational analysis, and effect on the collagen triple helix. Biopolymers 2008, 89, 443–454, eprint: https://onlinelibrary.wiley.com/doi/pdf/10.1002/bip.20864.

(10) Persikov, A. V.; Ramshaw, J. A. M.; Brodsky, B. Prediction of collagen stability from amino acid sequence. J. Biol. Chem. 2005, 280, 19343–19349.

(11) Cole, C. C.; Misiura, M.; Hulgan, S. A.; Peterson, C. M.; Williams III, J. W.; Kolomeisky, A. B.; Hartgerink, J. D. cation-π Interactions and Their Role in Assembling Collagen Triple Helices. Biomacromolecules 2022,

(12) Xu, F.; Zahid, S.; Silva, T.; Nanda, V. Computational design of a collagen A: B: C-type heterotrimer. J. Am. Chem. Soc. 2011, 133, 15260–15263.

(13) Persikov, A. V.; Ramshaw, J. A. M.; Kirkpatrick, A.; Brodsky, B. Amino Acid Propensities for the Collagen Triple-Helix. Biochemistry 2000, 39, 14960–14967, Publisher: American Chemical Society.

(14) Hentzen, N. B.; Islami, V.; Köhler, M.; Zenobi, R.; Wennemers, H. A Lateral Salt Bridge for the Specific Assembly of an ABC-Type Collagen Heterotrimer. J. Am. Chem. Soc. 2020, 142, 2208–2212, Publisher: American Chemical Society.

(15) Canty, E. G.; Kadler, K. E. Collagen fibril biosynthesis in tendon: a review and recent insights. *Comp. Biochem. Physiol.*, Part A: Mol. Integr. Physiol. 2002, 133, 979–985.

(16) Leikin, S.; Rau, D.; Parsegian, V. Temperature-favoured assembly of collagen is driven by hydrophilic not hydrophobic interactions. Nature structural biology 1995, 2, 205– 210.

(17) Anfinsen, C. B. Principles that govern the folding of protein chains. Science 1973, 181, 223–230.

(18) Stultz, C. M. The folding mechanism of collagen-like model peptides explored through detailed molecular simulations. Protein Sci. 2006, 15, 2166–2177.

(19) Baum, J.; Brodsky, B. Folding of peptide models of collagen and misfolding in disease. Curr. Opin. Struct. Biol. 1999, 9, 122–128.

(20) Park, S.; Klein, T. E.; Pande, V. S. Folding and misfolding of the collagen triple helix: Markov analysis of molecular dynamics simulations. Biophys. J. 2007, 93, 4108–4115.

(21) Hartmann, J.; Zacharias, M. Mechanism of collagen folding propagation studied by Molecular Dynamics simulations. *PLOS Comp*. Biol. 2021, 17, e1009079, Publisher: Public Library of Science.

(22) Bretscher, L. E.; Jenkins, C. L.; Taylor, K. M.; DeRider, M. L.; Raines, R. T. Conformational stability of collagen relies on a stereoelectronic effect. J. Am. Chem. Soc. 2001, 123, 777–778.

(23) Bachmann, A.; Kiefhaber, T.; Boudko, S.; Engel, J.; Bächinger, H. P. Collagen triple-helix formation in all-trans chains proceeds by a nucleation/growth mechanism with a purely entropic barrier. Proc. Natl. Acad. Sci. U. S. A. 2005, 102, 13897–13902.

(24) Favretto, F.; Flores, D.; Baker, J. D.; Strohäker, T.; Andreas, L. B.; Blair, L. J.; Becker, S.; Zweckstetter, M. Catalysis of proline isomerization and molecular chaperone activity in a tug-of-war. Nat. Comm. 2020, 11, 6046.

(25) Chen, J.; Edwards, S. A.; Grater, F.; Baldauf, C. On the cis to trans isomerization of prolyl–peptide bonds under tension. J. Phys. Chem. B 2012, 116, 9346–9351.

(26) Buevich, A. V.; Dai, Q.-H.; Liu, X.; Brodsky, B.; Baum, J. Site-Specific NMR Monitoring of cis-trans Isomerization in the Folding of the Proline-Rich Collagen Triple Helix. Biochemistry 2000, 39, 4299–4308.

(27) Baum, J.; Brodsky, B. Real-time NMR investigations of triple-helix folding and collagen folding diseases. Folding Des. 1997, 2, R53–R60.

(28) Tanrikulu, I. C.; Westler, W. M.; Ellison, A. J.; Markley, J. L.; Raines, R. T. Templated collagen “double helices” maintain their structure. J. Am. Chem. Soc. 2020, 142, 1137– 1141.

(29) Tanrikulu, I. C.; Raines, R. T. Optimal interstrand bridges for collagen-like biomaterials. J. Am. Chem. Soc. 2014, 136, 13490–13493.

(30) Fallas, J. A.; Lee, M. A.; Jalan, A. A.; Hartgerink, J. D. Rational Design of Single-Composition ABC Collagen Heterotrimers. J. Am. Chem. Soc. 2012, 134, 1430–1433.

(31) Zheng, H.; Lu, C.; Lan, J.; Fan, S.; Nanda, V.; Xu, F. How electrostatic networks modulate specificity and stability of collagen. Proc. Natl. Acad. Sci. U. S. A. 2018, 115, 6207–6212.

(32) Walker, D. R.; Hulgan, S. A. H.; Peterson, C. M.; Li, I.-C.; Gonzalez, K. J.; Hart-gerink, J. D. Predicting the stability of homotrimeric and heterotrimeric collagen helices. Nat. Chem. 2021, 13, 260–269.

(33) Madhan, B.; Xiao, J.; Thiagarajan, G.; Baum, J.; Brodsky, B. NMR monitoring of chain-specific stability in heterotrimeric collagen peptides. J. Am. Chem. Soc. 2008, 130, 13520–13521.

(34) Buevich, A.; Baum, J. Nuclear Magnetic Resonance Characterization of Peptide Models of Collagen-Folding Diseases. *Philos. Trans. R. Soc.*, B 2001, 356, 159–168, Publisher: The Royal Society.

(35) Saccà, B.; Renner, C.; Moroder, L. The chain register in heterotrimeric collagen peptides affects triple helix stability and folding kinetics. J. Mol. Biol. 2002, 324, 309–318.

(36) Matagne, A.; Radford, S. E.; Dobson, C. M. Fast and slow tracks in lysozyme folding: insight into the role of domains in the folding process. J. Mol. Biol. 1997, 267, 1068– 1074.

(37) Chaffotte, A. F.; Guillou, Y.; Goldberg, M. E. Kinetic resolution of peptide bond and side chain far-UV circular dichroism during the folding of hen egg white lysozyme. Biochemistry 1992, 31, 9694–9702.

(38) Simpson, R. B.; Kauzmann, W. The Kinetics of Protein Denaturation. I. The Behavior of the Optical Rotation of Ovalbumin in Urea Solutions1. J. Am. Chem. Soc. 1953, 75, 5139–5152.

(39) Greenfield, N. J. Analysis of the kinetics of folding of proteins and peptides using circular dichroism. Nature protocols 2006, 1, 2891–2899.

(40) Rigaku Oxford Diffraction, CrysAlisPro Software system.

(41) McCoy, A. J.; Grosse-Kunstleve, R. W.; Adams, P. D.; Winn, M. D.; Storoni, L. C.; Read, R. J. Phaser crystallographic software. J. Appl. Crystallogr 2007, 40, 658–674.

(42) Uśon, I.; Sheldrick, G. M. An introduction to experimental phasing of macromolecules illustrated by *SHELX*; new autotracing features. Acta Crystallogr. 2018, *D74*, 106–116.

(43) Caballero, I.; Sammito, M.; Millán, C.; Lebedev, A.; Soler, N.; Usón, I. ARCIMBOLDO on coiled coils. Acta Crystallogr. 2018, *D74*, 194–204.

(44) Fallas, J. A.; Dong, J.; Tao, Y. J.; Hartgerink, J. D. Structural Insights into Charge Pair Interactions in Triple Helical Collagen-like Proteins. J. Biol. Chem. 2012, 287, 8039–8047.

(45) Langer, G. G.; Cohen, S. X.; Lamzin, V. S.; Perrakis, A. Automated macromolecular model building for X-ray crystallography using ARP/ wARP version 7. Nature Protocols 2008, 3, 1171–1179.

(46) Emsley, P.; Lohkamp, B.; Scott, W. G.; Cowtan, K. Features and Development of Coot. Acta Crystallogr. 2010, D66, 486–501.

(47) Liebschner, D. et al. Macromolecular structure determination using X-rays, neutrons and electrons: recent developments in Phenix. Acta Crystallogr. 2019, *D75*, 861–877.

(48) Yennamalli, R.; Arangarasan, R.; Bryden, A.; Gleicher, M.; Phillips, G. N., Jr Using a commodity high-definition television for collaborative structural biology. J. Appl. Crystallogr. 2014, 47, 1153–1157.

(49) Morin, A.; Eisenbraun, B.; Key, J.; Sanschagrin, P. C.; Timony, M. A.; Ottaviano, M.; Sliz, P. Cutting edge: Collaboration gets the most out of software. eLife 2013, 2 .

(50) Wishart, D. S.; Bigam, C. G.; Yao, J.; Abildgaard, F.; Dyson, H. J.; Oldfield, E.; Markley, J. L.; Sykes, B. D. 1 H, 13 C and 15 N chemical shift referencing in biomolecular NMR. J. Biomol. NMR 1995, 6, 135–140.

(51) Misiura, M.; Shroff, R.; Thyer, R.; Kolomeisky, A. B. *DLPacker: deep learning for prediction of amino acid side chain conformations in proteins*; 2022; Vol. 90; pp 1278– 1290.

(52) Schrödinger, L.; DeLano, W. PyMOL. http://www.pymol.org/pymol.

## References

(1) Gauba, V.; Hartgerink, J. D. Self-Assembled Heterotrimeric Collagen Triple Helices Directed through Electrostatic Interactions. J. Am. Chem. Soc. 2007, 129, 2683–2690, Publisher: American Chemical Society.

(2) Buevich, A.; Baum, J. Nuclear Magnetic Resonance Characterization of Peptide Models of Collagen-Folding Diseases. *Philos. Trans. R. Soc.*, B 2001, 356, 159–168, Publisher: The Royal Society.

(3) Baum, J.; Brodsky, B. Folding of peptide models of collagen and misfolding in disease. Curr. Opin. Struct. Biol. 1999, 9, 122–128.

(4) Bachmann, A.; Kiefhaber, T.; Boudko, S.; Engel, J.; Bächinger, H. P. Collagen triple-helix formation in all-trans chains proceeds by a nucleation/growth mechanism with a purely entropic barrier. Proc. Natl. Acad. Sci. U. S. A. 2005, 102, 13897–13902.

(5) Kawahara, K.; Nemoto, N.; Motooka, D.; Nishi, Y.; Doi, M.; Uchiyama, S.; Nakazawa, T.; Nishiuchi, Y.; Yoshida, T.; Ohkubo, T.; Kobayashi, Y. Polymorphism of Collagen Triple Helix Revealed by 19F NMR of Model Peptide [Pro-4(R)-Hydroxyprolyl-Gly]3-[Pro-4(R)-Fluoroprolyl-Gly]-[Pro-4(R)-Hydroxyprolyl-Gly]3. J. Phys. Chem. B 2012, 116, 6908–6915, Publisher: American Chemical Society.

(6) Greenfield, N. J. Using circular dichroism collected as a function of temperature to determine the thermodynamics of protein unfolding and binding interactions. Nature protocols 2006, 1, 2527–2535.

(7) Hulgan, S. A.; Hartgerink, J. D. Recent Advances in Collagen Mimetic Peptide Structure and Design. Biomacromolecules 2022, 23, 1475–1489.

